# A mitochondrial SET domain protein is essential in *T. brucei* and required for mitochondrial morphology and function

**DOI:** 10.1101/2024.02.23.581798

**Authors:** Emily W. Knight, Rebeca Aquino Ventura, Niharikha Mukala, Meredith T. Morris

## Abstract

Protein lysine methylation is a ubiquitous post-translational modification. First identified in histone tails, many non-histone proteins are methylated at lysine residues. SET domain proteins catalyze the transfer of a methyl group from a donor such as S-adenosyl-L-methionine (SAM) to lysine residues. In this work, we identify a novel SET domain protein, TbSETD, in the African trypanosome, *Trypanosoma brucei*, and demonstrate that it is essential in both insect stage procyclic form and bloodstream form parasites. Epitope-tagged TbSETD localizes to the mitochondrion and TbSETD-deficient parasites exhibit growth defects, altered mitochondrial morphology, increased ROS levels, and increased sensitivity to apoptotic triggers. TbSETD binding proteins identified in immunoprecipitation experiments were enriched in mitoribosomal proteins. We also demonstrate that TbSETD immunoprecipitated from parasites has methyltransferase activity *in vitro* and methyllysine western blots reveal that TbSETD-deficient parasites have reduced levels of a 37 kDa mitochondrial protein. OrthoMCL DB indicates TbSETD is conserved only in kinetoplastids, suggesting a unique role in the biology of these parasites, and highlighting its potential as a drug target.

**Significance:** Kinetoplastid parasites include several medically important organisms for which current treatments are insufficient. SET domain protein lysine methyltransferases (PKMTs) are highly druggable and have predominately been studied in the context of chromatin remodeling and gene expression. This work provides the first example of a SET domain PKMT that localizes to the mitochondria in any protozoan parasite and the first implication that protein lysine methylation may be important to mitoribosomal biogenesis.

## Introduction

The flagellated protists in the group kinetoplastida include *Trypanosoma brucei, Trypanosoma cruzi,* and *Leishmania* spp. *T. brucei* and *T. cruzi* cause human African trypanosomiasis and Chagas disease, respectively, and *Leishmania* infections have multiple manifestations ranging from cutaneous leishmaniasis to more invasive visceral and mucocutaneous infections (1). There has been a recent increase in human cutaneous leishmaniasis in the United States (2, 3). The current treatments for these diseases are sub-optimal and the development of drug resistance necessitates the search for additional drug targets.

Protein methylation is a ubiquitous and dynamic post-translational modification that alters protein activity, half-life, localization, and protein-protein interactions. Methylation can occur on lysine, arginine, and histidine residues. The best studied of these modifications in kinetoplastids is arginine methylation, a modification found on RNA binding proteins that plays a role in the gene expression regulation (4). Silencing the protein arginine methyltransferase PRMT1 in *T. brucei* had wide-ranging impacts, including leading to increased abundance of glycolytic enzymes and impaired RNA granule formation in response to stress (5). These findings highlight the importance of non-histone protein methylation as a mechanism to regulate trypanosome cell biology.

SET domain protein lysine methyltransferases (PKMTs) catalyze the transfer of methyl groups from S-adenosyl methionine to lysine resides in proteins. First recognized for modifying histones, it is now clear that SET domain PKMTs can catalyze methylation of many non-histone substrates including tumor suppressors, proteins involved in signaling pathways and translation, and transcription factors (6). In the single study of protein lysine methylation in trypanosomes, methylated lysine residues were identified in proteins related to translation, stress response, glycolytic processes, amino acid transport, and cellular metabolism (7) . The role that SET domain PKMTs play in kinetoplastid biology, especially through the modification of non-histone proteins, is unknown.

The trypanosome genome encodes 40 proteins that contain SET domains. Thirty of these proteins were annotated as SET domain proteins with homologs in other organisms (8) while ten are annotated as hypothetical or conserved with no identifiable orthologs in other organisms. We are interested in defining the function of these ten unique SET domain proteins in parasite biology.

Here we demonstrate that Tb927.9.11350 (TbSETD), which is annotated as a hypothetical protein containing a SET domain, is essential in both insect stage procyclic form (PF) and mammalian bloodstream form (BF) parasites. In PF parasites, the protein localized to the mitochondrion and PF TbSETD-deficient parasites had altered mitochondrial morphology. Depletion of TbSETD in either PF or BF parasites led to increased levels of intracellular reactive oxygen species. Interestingly, PF TbSETD-deficient parasites were more sensitive to apoptotic triggers while BF TbSETD-deficient parasites were not. Proteins identified from TbSETD immunoprecipitations were enriched in mitoribosome components. Furthermore, TbSETD immunoprecipitated from parasites had methyltransferase activity *in vitro* and methylation of a mitochondrial 37kDa protein was reduced in PF cells lacking TbSETD. Taken together, these observations indicate that TbSETD is an essential mitochondrial SET domain methyltransferase that is required for normal mitochondrial function and suggests a potential role in mitoribosome assembly and mitochondrial gene regulation.

## Methods

### Cell culture and transfection of *T. brucei*

Procyclic form (PF) 29-13 and bloodstream form (BF) 90-13 parasites expressing T7 polymerase and tetracycline (tet) repressor (9) were maintained in SDM79 (or the minimal glucose version SDM79θ containing 5 μM glucose) (10, 11) and HMI-9 culture medium (12), respectively. The plasmid vectors for the expression of epitope tagged TbSETD were generated by fusing the V5 epitope sequence to the 5’ end of the TbSETD open reading frame or the hemagglutinin antigen sequence to the 3’ end of the TbSETD open reading frame. Both fusions were cloned into the pXS2 (PF) expression vector possessing a blasticidin resistance gene (13). For transfection, 20 μg of plasmid DNA was linearized by restriction enzyme treatment (pXS2 and pXS6 plasmids: MluI; pZJM plasmids: NotI) and electroporated in 4 mm cuvettes (BioRad GenePulser Xcell; exponential, 1.5 kV, 25 μF). Twenty-four hours after electroporation, culture media was supplemented with the appropriate drug for selection: 15 μg/ml G418; 50 μg/ml hygromycin; 2.5 μg/ml phleomycin; 10 μg/ml blasticidin. RNA interference (RNAi) cell lines were generated by cloning nucleotides 60-175 of TbSET into the inducible pZJM vector to generate the plasmid pZJMTbSETD.Phleo. BF 90-13 TbSETD RNAi cell lines could not be recovered from frozen stocks. Consequently, cells were transformed prior to each experiment.

### Immunofluorescence microscopy

All steps are performed at room temperature (RT). Cells were harvested (800 x g, 10 min), washed once with PBS, incubated with 150 nM MitoTracker Red CMXRos (Invitrogen, M7512) for 30 min, washed twice with PBS, fixed with 2% paraformaldehyde in PBS for 30 min and allowed to settle on slides for 45 min. Adhered cells were washed once with wash solution (0.1% normal goat serum in PBS) and permeabilized with 0.5% Triton X-100 for 30 min. Following permeabilization, cells were washed twice with wash solution and blocked with 10% normal goat serum (NGS) in PBS with 0.1% Triton X-100. Primary antibodies (monoclonal V5, Thermo Scientific, 1:500) were diluted in block solution and incubated with cells for 1 h. Following primary antibody incubation, slides were washed five times with wash solution and incubated with secondary antibody (goat anti-Alexa fluor 488, Thermo Fisher, 1:1,000) diluted in block solution for 1 h. Slides were then washed another five times with wash solution, mounted using VECTASHIELD Mounting Medium (Vector), and imaged using a Leica DMi8 microscope.

### Growth curves

PF cells expressing HA-tagged TbSETD and possessing the RNAi inducible pZJMTbSETD vector (PF TbSETD.HA.KD and BF TbSETD.HA.KD) were seeded at 1 x 10^5^ cells/mL in SDM79 or SDM79θ (PF) or 5 x 10^4^ cells/mL HMI-9 (BF) and RNAi induced with 1 μg/ml doxycycline. PF cells were allowed to grow to a density of 5 x 10^6^ cells/mL prior to passing to 1 x 10^5^ cells/mL. BF cells were allowed to grow to a density of 1 x 10^6^ cells/mL prior to passing back to 5 x 10^4^ cells/mL. Culture density was monitored by flow cytometry at 24 h intervals using a CytoFLEX (Beckman Coulter).

### Electron microscopy processing and imaging

PF Cells were harvested (5 x 10^7^, 800 x g, 10 min), washed three times with PBS, and fixed (2% paraformaldehyde, 2.5% glutaraldehyde in 100 mM phosphate buffer pH 7.4). Cells were stored at 4°C for no more than two days before being processed as described previously (14).

### Immunoprecipitations (IPs) for mass spectrometry

All steps were performed at 4° C. PF cells expressing HA-tagged TbSETD (10⁹; PF TbSETD.HA) were harvested (800 x g, 10 min) and washed twice with PBS. The cells were lysed in hypotonic lysis buffer 1 (HB1) (100 μM tosyllysine chloromethyl ketone [TLCK], 1 μg/mL leupeptin in H_2_O) and incubated on ice for 20 min. An equal volume of hypotonic lysis buffer 2 (HB2) was added (100 mM Hepes-KOH pH 8, 50 mM KCl, 10 mM MgCl_2_, 20% (v/v) glycerol, 100 μm TLCK, 1 μg/mL leupeptin). The lysates were then passed through a 28-guage needle three times and centrifuged (17,000 x g, 10 min, 4° C), and lysate (L1) was transferred to a new tube. The pellet was resuspended in equal volumes of HB1 and HB2 supplemented with 0.5% IGEPAL and 1X phosphatase HALT inhibitor cocktail (Thermo Fisher) and incubated on ice for 20 min. The lysate from the resuspended pellet was then centrifuged (17,000 x g, 10 min, 4° C) and lysate (L2) was transferred to a new tube. Magnetic anti-HA beads (Thermo Scientific) were added to the L2 lysate and incubated, rotating end-over-end overnight at 4° C. The beads were separated using a magnetic rack, washed three times with 5X TBS-T (100 mM Tris pH 7.5, 750 mM NaCl, 0.5% Tween20), washed twice with nanopure water, and resuspended in 100 μl nanopure water. Samples were then analyzed by mass spectrometry as described below.

### LC-MS/MS analysis of IPs

IPs were run on NuPAGE Bis-Tris 4-12% gradient gel (ThermoFisher). Each SDS-PAGE gel lane was sectioned into 12 segments of equal volume. Each segment was subjected to in-gel trypsin digestion as follows. Gel slices were destained in 50% methanol (Fisher), 100 mM ammonium bicarbonate (Sigma-Aldrich), followed by reduction in 10 mM Tris[2-carboxyethyl]phosphine (Pierce) and alkylation in 50 mM iodoacetamide (Sigma-Aldrich). Gel slices were then dehydrated in acetonitrile (Fisher), followed by addition of 100 ng porcine sequencing grade modified trypsin (Promega) in 100 mM ammonium bicarbonate (Sigma-Aldrich) and incubation at 37°C for 12-16 hours. Peptide products were then acidified in 0.1% formic acid (Pierce). Tryptic peptides were separated by reverse phase XSelect CSH C18 2.5 μm resin (Waters) on an in-line 150 x 0.075 mm column using a nanoAcquity UPLC system (Waters). Peptides were eluted using a 30 min gradient from 97:3 to 67:33 buffer A:B ratio. [Buffer A = 0.1% formic acid, 0.5% acetonitrile; buffer B = 0.1% formic acid, 99.9% acetonitrile.] Eluted peptides were ionized by electrospray (2.15 kV) followed by MS/MS analysis using higher-energy collisional dissociation (HCD) on an Orbitrap Fusion Tribrid mass spectrometer (Thermo) in top-speed data-dependent mode. MS data were acquired using the FTMS analyzer in profile mode at a resolution of 240,000 over a range of 375 to 1500 m/z. Following HCD activation, MS/MS data were acquired using the ion trap analyzer in centroid mode and normal mass range with precursor mass-dependent normalized collision energy between 28.0 and 31.0. Proteins and post-translational modifications were identified by database search using Mascot (Matrix Science) with a parent ion tolerance of 3 ppm and a fragment ion tolerance of 0.5 Da.

Scaffold (Proteome Software) was used to verify MS/MS based peptide and protein identifications. Peptide identifications were accepted if they could be established with less than 1.0% false discovery by the Scaffold Local FDR algorithm. Protein identifications were accepted if they could be established with less than 1.0% false discovery and contained at least 2 identified peptides.

### Immunoprecipitations (IPs) for Activity Assays

All steps were performed at 4° C. PF cells expressing HA-tagged TbSETD (10⁹; PF TbSETD.KD) were harvested (800 x g, 10 min) and washed twice with PBS. The cells were lysed in hypotonic lysis buffer 1 (HB1) (100 μM tosyllysine chloromethyl ketone [TLCK], 1 μg/mL leupeptin) and an equal volume of hypotonic lysis buffer 2 (HB2) (100 mM Hepes-KOH pH 8, 50 mM KCl, 10 mM MgCl_2_, 20% (v/v) glycerol, 100 μm TLCK, 1 μg/mL leupeptin). The mixture was supplemented with 0.5% IGEPAL and incubated on ice for 20 min. Magnetic anti-HA beads (Thermo Scientific) were added to the mixture and incubated, rotating end-over-end overnight at 4° C. The beads were separated using a magnetic rack, washed five times with 5X TBS-T (100 mM Tris pH 7.5, 750 mM NaCl, 0.5% Tween20), washed twice with nanopure water, and resuspended in 50 μl nanopure water. Samples were then analyzed by Western Blotting and Methylation Activity Assays as described below.

### ROS Measurement

PF and BF form *T. brucei* (5 x 10^6^ cells) were centrifuged at 800 x g for 10 minutes. Cells were resuspended with 1mL PBS+ 0.1% glucose (PBSg). Samples (500 μL) were treated with 1 μL ROS stain (Total Reactive Oxygen Species (ROS) Assay Kit 520nm, Invitrogen) and the other 500 μL with 1 μL DMSO. Cells were incubated for 60 min at either 25°C and 5% CO_2_ (PF) or 37°C and 5% CO_2_ (BF). The samples were split into 250 μL each, treated with either 125 μM H_2_O_2_ or equal volume PBSg, incubated at 25°C and 5% CO_2_ (PF) or 37°C and 5% CO_2_ (BF) for 60 min and analyzed using a CytoFLEX flow cytometer (Beckman Coulter). Events (10,000) were recorded and analyzed using either FlowJo software or FCSexpress software. Each experiment was performed with biological triplicates in technical triplicates.

### Apoptosis assay using AnnexinV-FITC

PF or BF form *T. brucei* (5 x 10^6^) were pelleted by centrifugation at 800 x g for 10 min. Cells were then resuspended in fresh media (SDM79 or SDM798 for PF and HMI-9 for BF) containing 2 mM H_2_O_2_(15) and incubated for 180 min at 25°C and 5% CO_2_ for PF and 37°C and 5% CO_2_ for BF. The cells were washed once with 1X PBS, resuspended in 200 μL binding buffer (10 mM HEPES/NaOH, pH 7.4, 140 mM sodium chloride, 2.5mM CaCl_2_) and 2 μL Annexin V-FITC and 2 μL propidium iodide (Enzo Life Sciences) and incubated for 15 min. Cells were then analyzed using a CytoFLEX flow cytometer (Beckman Coulter). Values for FITC-H and PC5.5-H were recorded for 10,000 events and analyzed using FCSexpress software. Each experiment was performed with biological triplicates in technical triplicates. The data are expressed as a percentage of the Annexin V-FITC positive cells in the total population. Median fluorescence intensity (MFI) was calculated, and Welch’s T-tests were performed (ns: not statistically significant, *P < 0.05, **P < 0.01, ***P < 0.001).

### Quantitation of glycosome and mitochondrial size via TEM

Measurements of glycosome size and cell area in TEM images were performed using FIJI. Organellar area was calculated by outlining single glycosomes or mitochondria and measuring image area. To calculate organellar area as a percentage of total cell area, the sum area of measured organelles was divided by the total cell area measured. Welch’s T-tests were performed (ns: not statistically significant, *P < 0.05, **P < 0.01, ***P < 0.001).

### Methyltransferase assays

Assays were performed using Promega MTase-Glo Methyltransferase Assay (Promega V7601) with the following modifications. The protein-magnetic bead IP slurry was diluted 1:1 in 1x reaction buffer (20 mM Tris buffer (pH 8.0), 50 mM NaCl, 1 mM EDTA, 3 mM MgCl2, 0.1 mg/ml BSA, and 1 mM dithiothreitol (DDT)). Following the initial dilution, a two-fold serial dilution with 1X reaction buffer was conducted in clear PCR tubes (VWR). The methyltransferase reaction was initiated by adding 10 ml of 2x substrate solution (80 mM Tris pH 8.0, 200mM NaCl, 1 mM EDTA, 3 mM MgCl_2_, 0.4 mg/ml BSA and 1 mM DTT, 1mM S-adenosyl methionine (SAM), 10μM calf thymus histones (Roche, catalog number 10223565001) and nanopure water to each tube. The reactions were then incubated at 37 °C with continuous shaking for 2 h on an orbital shaker. After incubation, the magnetic beads were separated from the reactions by positioning the tubes near a magnet and transferring the reactions to a 96-well clear plate (BRANDplates®, Germany). After bead removal, the conversion of produced S-Adenosyl-L-homocysteine (SAH) to Adenosine diphosphate (ADP) was initiated by supplementing each reaction in the assay plate with 5x MTase-Glo Reagent (prepared by diluting 10x MTase-Glo reagent with nanopure water). The plate was then mixed for 2 min and incubated 30 min at room temperature. MTase-Glo detection solution was added to the reaction mixture, the plate was mixed for 2 min incubated for 30 min at room temperature. Finally, the released luminescence was quantified using the Synergy™ H1 luminometer (BioTek, USA).

### Methyllysine Western blotting

All steps were performed at 4° C. PF Cells expressing HA-tagged TbSETD (PF TbSETD.HA) or HA-tagged TbSETD cells possessing the RNAi inducible pZJM TbSETD vector (PF TbSETD.HA.KD). Knockdown cells were induced with 1 μg/ml doxycycline for either 24 or 48 h. For each cell line, 10^9^ cells were harvested (800 x g, 10 min), washed twice with PBS, and lysed using a 1 volume wet-weight silicon carbide abrasive (1:1 silicon carbide combined with STE [250mM Sucrose, 25 mM Tris-HCl pH 7.4, 1 mM EDTA] with a protease inhibitor tablet [Pierce Protease Inhibitor Mini Tablets] and 1 mM PMSF). Breakage was confirmed by microscopy and abrasive removed by centrifugation (100 x g, 1 min), the supernatant transferred to a new tube. Nuclei were pelleted (2,000 x g, 10 min) and the supernatant transferred to a new tube. Mitochondria were pelleted for (5,000 x g, 10 min), resuspended in STE, protein concentration measured via BCA. Protein (20-30 μg) was precipitated (16) with 100 mM NaCl, 4X volumes of acetone and incubated for 30 min at room temperature after mixing. Samples were centrifuged at room temperature (17,000 x g, 10 min), the supernatant was removed, and the pellet dried (open air benchtop until acetone evaporated). SDS (20 μL of 1%) was added to the pellet, incubated at room temperature for 1 hr and sonicated for 30 minutes. Cracking Buffer (5 μL OF 3X) was added to the samples, which were separated by SDS-PAGE (4–15% Mini-PROTEAN® TGX™ Precast Protein Gels, Bio-Rad) and analyzed by Western blotting (Rab anti-KMe 1:2,500, abcam ab23366).

## Results

### T. brucei contains a novel SET domain protein

SET domain protein lysine methyltransferases (PKMTs) catalyze the post-translational addition of methyl groups to lysine residues of target proteins (17) which influences enzyme activity, protein:protein interactions, protein stability, and protein localization. To identify potential SET domain PKMTs, we queried TriTrypDB (https://tritrypdb.org) with the keyword “SET” and identified 43 proteins (**Table 1**). For 40 of these proteins, AlphaFold predictions and InterPro domains confirmed the presence of a SET domain. We were unable to identify SET domains in three of the genes annotated as SET domain proteins (**Table 1**, listed as “No data available”). In the list of 43 putative SET PKMTs, ten open reading frames were hypothetical or conserved unknown, thirty had homologs in other eukaryotes (8) two were annotated as polyadenylate-binding proteins 1 and 2, and one was annotated as a putative NADH dehydrogenase (ubiquinone) 1 beta subcomplex subunit. Except for SET26 and SET27, which play a role in transcription initiation and termination (8) the role that SET domain PKMTs play in the regulation of parasite biology via lysine methylation is unknown.

**Table 1.**
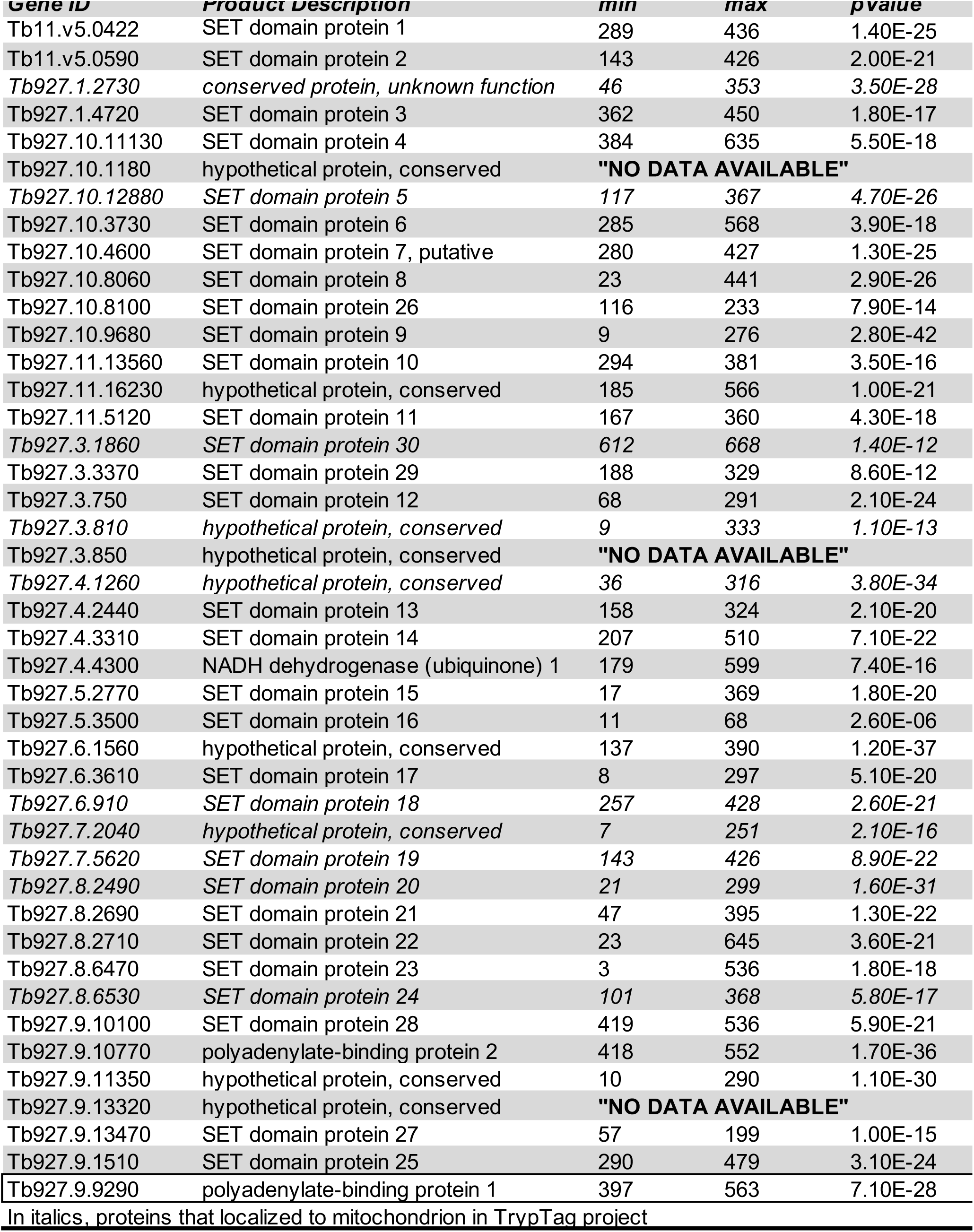
SET domain proteins in *T. brucei* 927 genome

In this work, we focused on Tb927.9.11350. OrthoMCL DB (https://orthomcl.org) indicated this protein was restricted to kinetoplastids including *Leishmania*, *T. brucei* and *T. cruzi*. SET domains are ∼ 130 residues long and characterized by a general three-dimensional structure containing a series of three sheets of β-structures folding around a pseudo-knot structure (17). SET domain proteins are poorly conserved at the primary amino acid level and Tb927.9.11350 was ∼ 20% identical to the closest SET protein we could identify, HsSETD6 **(Fig 1A**). Most SET proteins contain multiple domains and HsSETD6 is typical in that it contains an N-terminal helix, a SET domain, an i-SET domain, and a C-SET domain (18). In contrast, Tb927.9.11350 is a predicted 310 residue protein containing only a putative SET domain. To build confidence that Tb927.9.11350 contains a SET domain, we aligned the predicted AlphaFold structure (Q38DP6) with HsSETD6 (3QXY) (**Fig. 1B**). Overall, Tb927.9.11350 aligned well with the SET domain HsSETD6 with both structures sharing several conserved beta structures and alpha helices. While there are some similarities in their secondary structure, Tb927.9.11350 contains a C-terminal helix that is absent in HsSETD6 and Tb927.9.11350 lacks any identifiable functional domains outside of the SET domain. In multiple sequence alignments of other closely related SET domain proteins, the only invariant residue required for activity is Y261 (HsSETD6) (18). While there is a Y270 in Tb927.9.11350, it is unknown whether this residue is required for activity. Based on the structural similarities with HsSETD6 and the annotation in TritrypDB, we have named this unique SET protein TbSETD.

**Figure 1.**
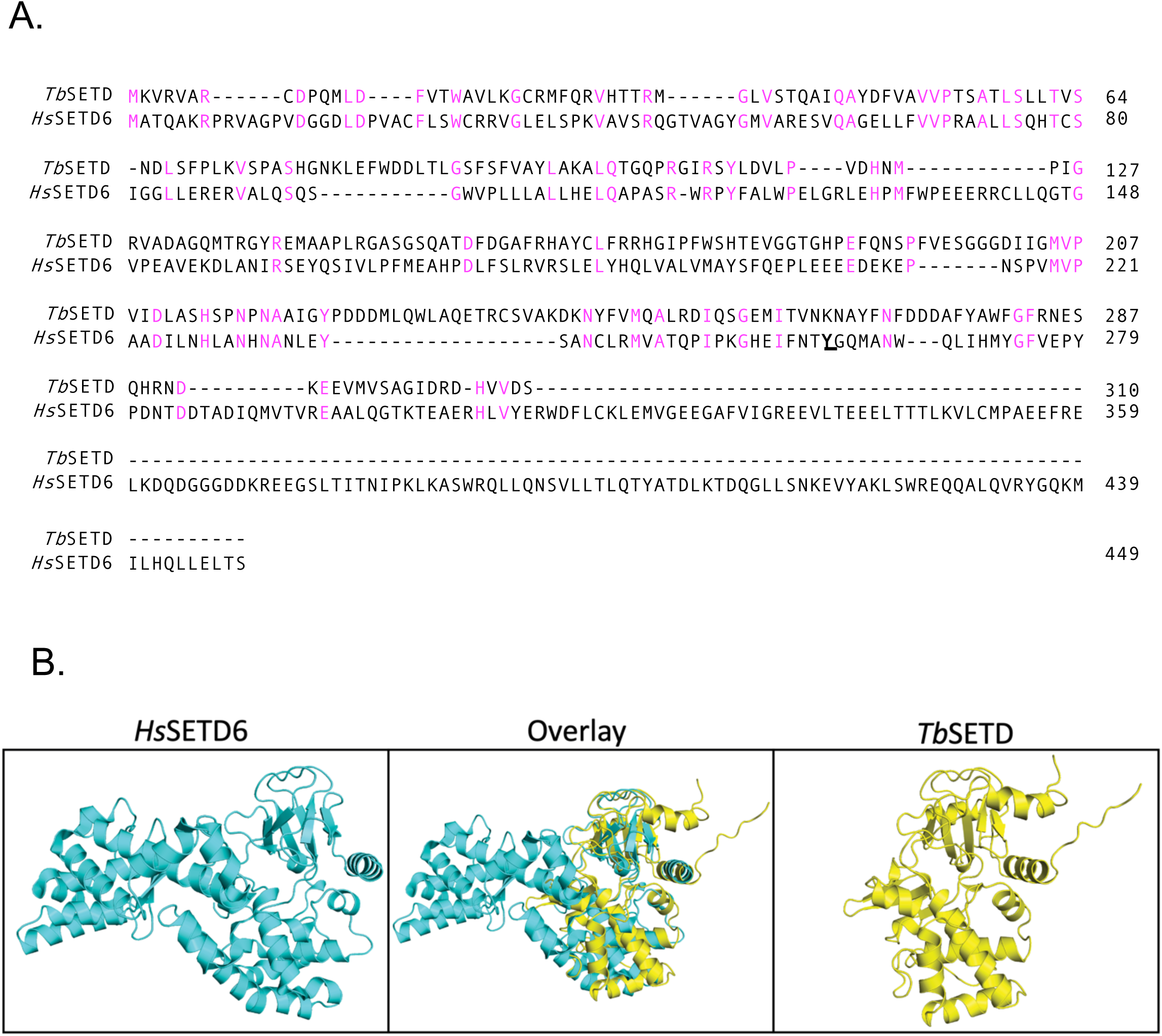
TbSETD contains a SET domain. **A.** Sequences for TbSETD (Tb927.9.11350, PDB: Q38DP6) and HsSETD6 (PDB: 3QXY) were aligned using Constraint-based Multiple Alignment Tool (https://www.ncbi.nlm.nih.gov/tools/cobalt/re_cobalt.cgi). Magenta amino acids are conserved between the two proteins. Y261 required for catalytic activity of HsSETD6 is underlined. **B**. PyMOLstructure of SETD6 in cyan was overlayed with the predicted AlphaFold structure of TbSETD in yellow.

### TbSETD localizes to mitochondria in PF parasites

Because the biological function of PKMTs is mediated through the modification of target substrates that localize with the protein, we wanted to determine the localization of TbSETD. To this end, we analyzed PF cell lines expressing TbSETD fused to an N-terminal V5 tag (PF V5.TbSETD). PF *T. brucei* has a single-branched mitochondrion that spans the length of the parasite. Anti-V5 antibody staining followed this pattern and overlapped with MitoTracker Red, which utilizes the mitochondrial membrane potential to stain mitochondria of live cells (**Fig. 2**). These images indicate that TbSETD is localized to the mitochondrion in PF parasites. Currently, TbSETD is not represented in the TrypTAG protein localization database (http://tryptag.org). Sequence analysis using PSORTb v.3.0 (19) (https://www.psort.org) and WoLF PSORT (https://wolfpsort.hgc.jp/) failed to identify any targeting sequences. However, a separate prediction method developed in *T. brucei* employing additional parameters such as physiochemical properties, secondary structure, unfoldability, and estimated radius of gyration predicted TbSETD as a mitochondrial protein (20).

**Figure 2.**
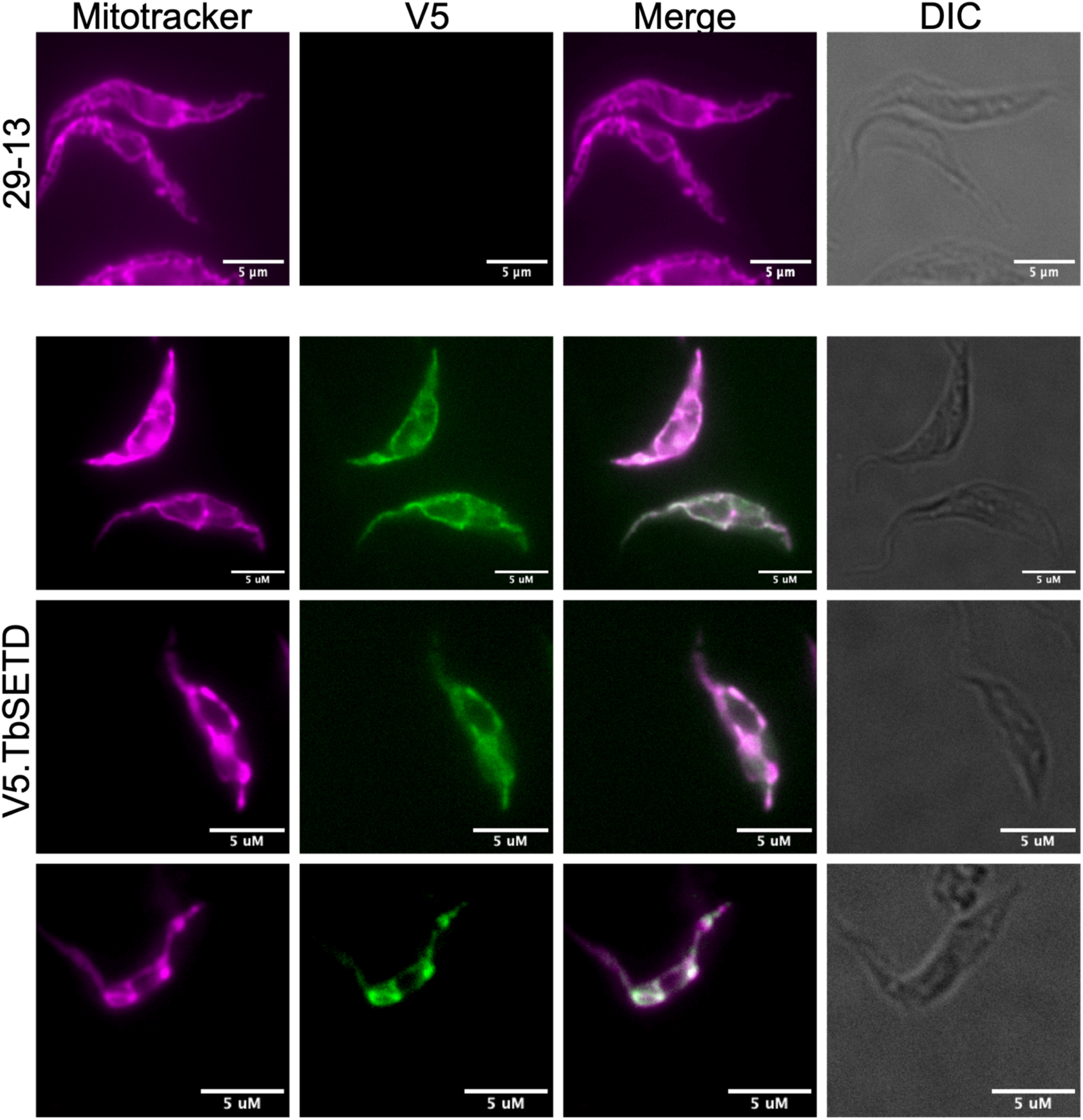
V5.TbSETD localizes to mitochondria. Control PF 29-13 parental cells and PF 29-13 cells expressing V5.TbSETD were stained with 150 nM MitoTracker (magenta), fixed, and labeled with antibodies against V5 (green). The V5 epitope was detected using AlexaFluor 488.

### TbSETD is essential in both PF and BF parasites

To assess the effect of TbSETD depletion on parasite biology, we next silenced TbSETD in both PF and BF parasites using RNA interference (RNAi). Because we do not have antibodies against recombinant TbSETD, knockdowns in PF cells were performed in cell lines expressing TbSETD fused to a C-terminal HA tag (PF TbSETD.HA.KD) so that we could monitor the penetrance of RNAi. In the fly, the parasites reside predominately in a glucose-deplete environment (21). However, in the laboratory, PF are routinely cultured in high-glucose media. To understand the impact of the loss of this protein in both conditions, we measured growth rates of PF TbSETD.HA.KD parasites grown in high-glucose (5 mM) and glucose-deplete (5 μM) conditions. By 72 hours, TbSETD.HA expression in PF parasites grown in either 5 μM or 5 mM glucose was reduced more than 90%. Silencing TbSETD also impacted growth, particularly in 5 μM glucose (**Fig. 3A**). In the glucose deplete medium, PF cells had a doubling time of ∼ 17 h while TbSET.HA.KD cells had a doubling time greater than 80 h. When the same cells were grown in glucose replete (5 mM) medium, no growth defect was detected (**Fig. 3B**).

**Figure 3.**
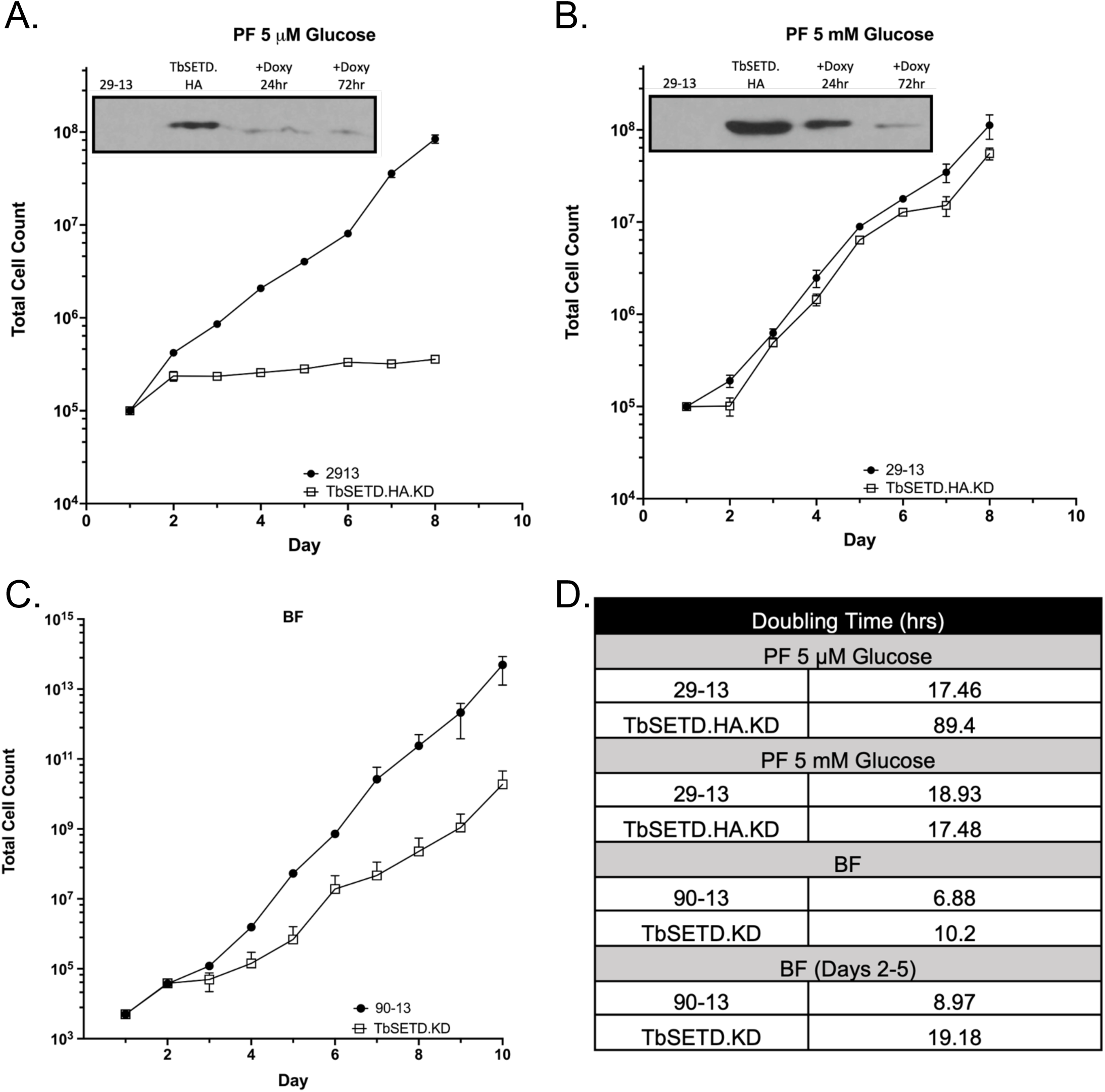
TbSETD.HA.KD cells exhibit slow growth rates. PF parasites expressing TbSETD.HA and harboring pZJMTbSETD construct for gene silencing and BF parasites harboring pZJMTbSETD for gene silencing were induced with doxycycline and counted daily. **A.** PF parasites grown in 5 µM glucose **B.** PF parasites grown in 5 mM glucose **C.** BF parasites **D.** Doubling times estimates. **Panel A and B insets**: Western blots of cell lysates probed with antibodies against the HA tag.

To assess the essentiality of this protein in BF parasites, we measured growth rates of parasites after RNAi in TbSETD.KD cells. In BF parasites, attempts to tag TbSETD were unsuccessful, so we could not follow changes in expression levels that resulted from silencing. Nevertheless, RNAi of TbSETD led to a marked growth phenotype when compared to parental cells (**Fig. 3C-D**). The growth defect was most pronounced on days 2-5 when BF TbSETD.KD cells had a doubling time of ∼19 h compared to that of parental 90-13 BF cells (∼9 h). After day 6, the growth rate of RNAi cell lines increased, likely because the cells had become refractory to silencing of essential proteins (22, 23). These results indicate that TbSETD is essential in BF parasites and PF parasites grown in low glucose and suggests that TbSETD activity may be less important when PF are grown in high glucose.

### PF TbSETD.HA.KD have altered mitochondria

Because TbSETD localizes to mitochondria and silencing the protein inhibited parasite growth, we used transmission electron microscopy (TEM) to determine how depletion of TbSETD altered PF mitochondrial morphology in both 5 μM and 5 mM glucose media (**Fig. 4**). In parental cells, mitochondria are present as extended tubular structures with a uniform diameter that spans much of the length of the cell body. In contrast, mitochondria in PF TbSETD.HA.KD cells were clearly abnormal, lacked this elongated structure, and exhibited a more rounded, spherical morphology (**Fig. 4A-B**). In high glucose conditions, mitochondrion cells occupied 0.12 ± 0.05 % and 0.16 ± 0.05 % of the cell area of PF 29-13 and TbSETD.HA.KD cells, respectively (**Fig. 4C**). This increased mitochondrial abundance was also observed in cells cultured in glucose deplete media, where mitochondria comprised 0.13 ± 0.04 and 0.19 ± 0.11 % of the cell area of PF 29-13 cells and TbSETD.HA.KD cells (**Fig. 4D**), respectively. In both cases, the difference in mitochondrial area was reproducible but not statistically significant. We also noted that the variation in mitochondrial area was highest in PF TbSETD.HA.KD grown in low glucose and ranged from less than 0.05 to greater than 0.3 percent of the cell area.

**Figure 4:**
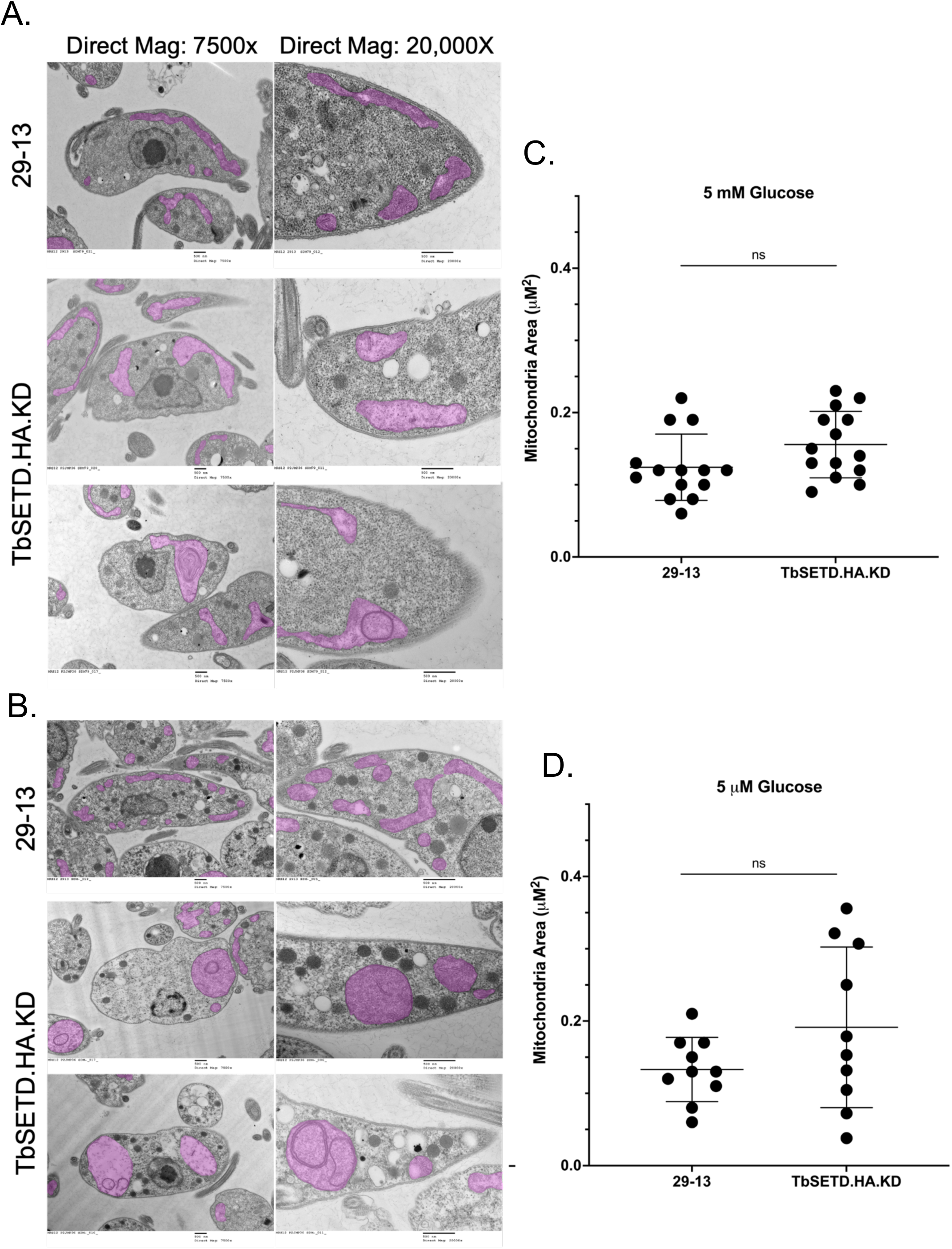
TbSETD.HA.KD cells have swollen mitochondria. TEM of PF parasites. **A.** TbSETD.HA.KD cells grown in 5 mM glucose **B.** TbSETD.HA.KD cells grown in 5 μM glucose **C.** Quantification of mitochondrial area of images in panel A represented as percent area of the cell **D.** Quantification of mitochondrial area of images in panel C represented as percent area of cell. Mitochondria are pseudocolored magenta.

To get another measurement of mitochondrial morphology changes in PF cells, we labeled mitochondria with MitoTracker Red. Mitochondria of PF 29-13 cells grown in 5 mM glucose were typical of “normal” trypanosomes, being tubular with a uniform branching pattern that spanned the length of the parasite (**Fig. 5A**). PF TbSETD.HA cells had mitochondria with increased branching. These cells were also shorter and thicker compared to parental 29-13 parasites. In contrast, PF TbSETD.KD had mitochondria with decreased branching and regions where the mitochondrion appeared to be thicker (**Fig. 5A, arrows**).

**Figure 5.**
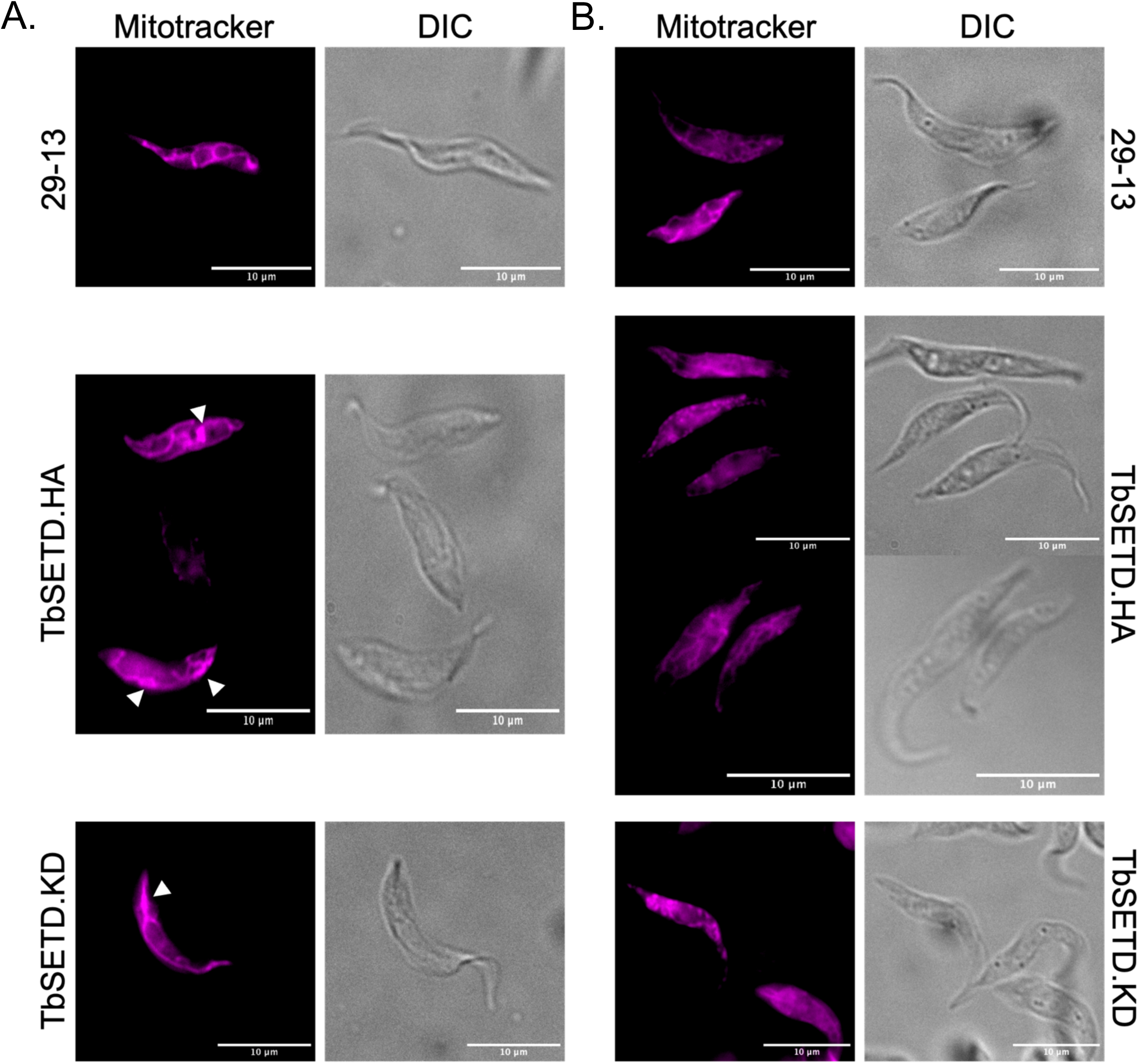
MitoTracker staining of TbSETD.HA and TbSETD.KD cells in 5 mM and 5 mM glucose. PF cells were stained with 150 nM MitoTracker Red, fixed, and imaged. **A.** Widefield fluorescence images of cells grown in 5 mM glucose. Arrows indicate areas of mitochondrial swelling **B.** Widefield fluorescence images of cells grown in 5 μM glucose

Mitochondria of parental PF 29-13 cells grown in 5 μM glucose are more intricate with a higher degree of branching in comparison to cells grown in 5 mM glucose (**Fig. 5B**). Under these conditions (low glucose), it was difficult to identify discrete portions of the mitochondria using MitoTracker Red staining. This prevented meaningful comparisons of the mitochondria from TbSETD.HA and TbSETD.KD cells with parental lines under the glucose replete condition.

### PF TbSETD-deficient cells grown in 5 μM glucose have smaller glycosomes

In other eukaryotes, mitochondria and peroxisome functions are coordinated and defects in one organelle often affect the other (24, 25). To assess the effect of TbSETD depletion on glycosomes, we measured glycosome size and number in PF TbSETD.KD cell lines grown in 5 μM and 5 mM glucose (**Fig. 6**). In 5 mM glucose media, the size difference of glycosomes was not statistically different between the two cell lines (**Fig. 6A**). Average glycosome area for PF 29-13 was 0.06 ± 0.02 μM^2^ while TbSETD.HA.KD was 0.08 ± 0.03 μM^2^ . In contrast, PF TbSETD.KD cells grown in 5 μM glucose media had 25% smaller glycosomes (0.06 ± 0.02 μM^2^) than the parental line (0.08 ± 0.02 μM^2^) (**Fig. 6B**). There was no difference in number of glycosomes between the two cell lines. In 5 mM glucose, parental cells had 54.88 ± 28.02 glycosomes/100mm^2^ while TbSETD.HA.KD had 65.80 ± 35.36 glycosomes/100mm^2^ (**Fig. 6C**). In low glucose, parental cells had 114.72 ± 68.64 glycosomes /100mm^2^ while TbSETD.HA.KD had 110.83 ± 52.36 glycosomes /100mm^2^ (**Fig. 6D**).

**Figure 6.**
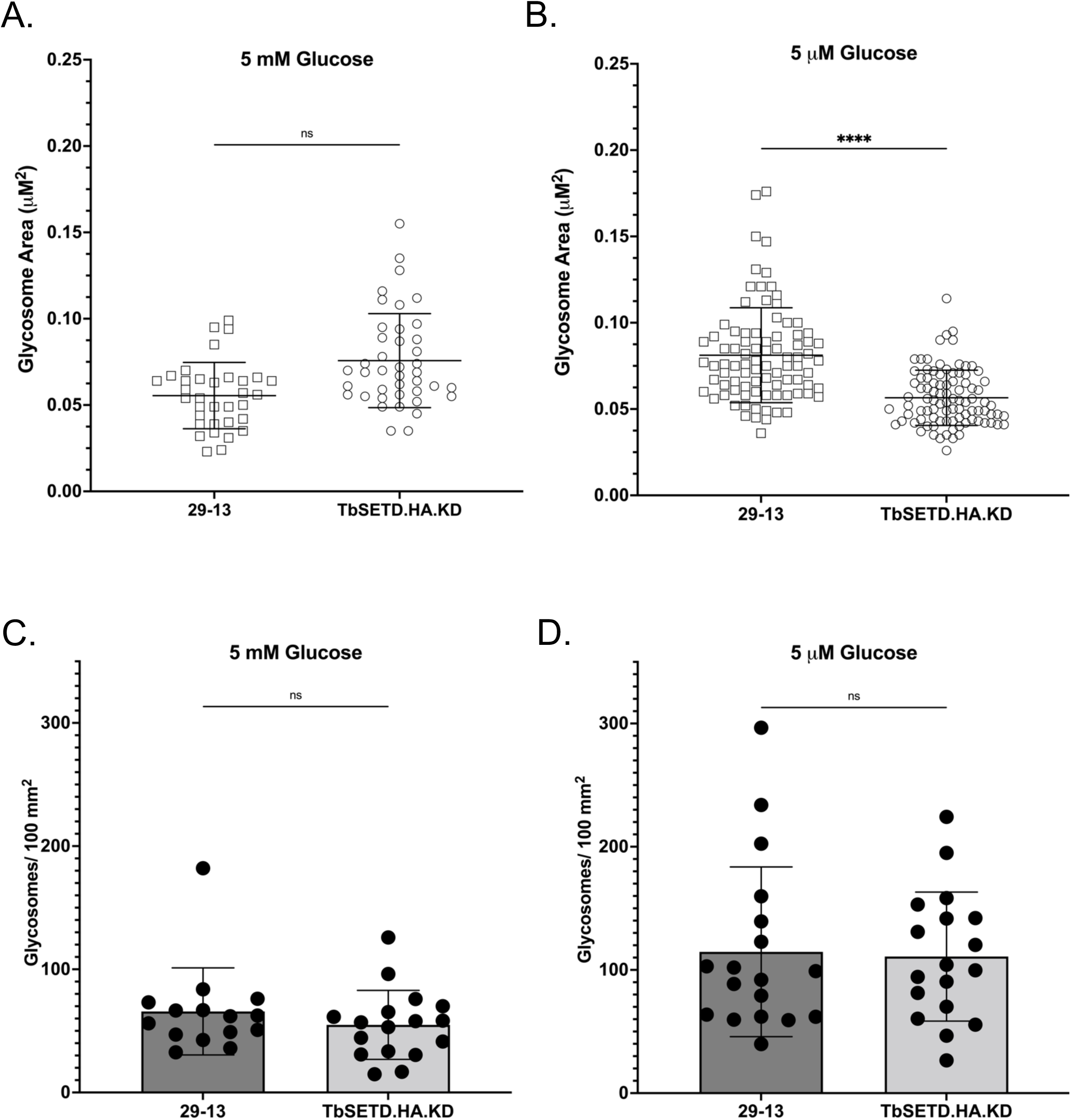
Glycosome size, but not number are altered in TbSETD.HA.KD cells. **A.** Bar graphs of glycosome area (μM^2^) of PF grown in 5 mM glucose. **B.** Bar graphs of glycosome area (μM^2^) of PF grown in 5 μM glucose. **C.** Bar graphs of glycosome number/100 mm^2^ of cells grown in 5 mM glucose **D.** Bar graphs of glycosome number/100 mm^2^ of cells grown in 5 μM glucose. Averages were calculated, and Welch’s T-tests were performed (*P < 0.05, **P < 0.01, ***P < 0.001).

### BF TbSETD.KD and PF TbSETD.KD cells had increased ROS, and PF TbSETD.KD but not BF TbSETD.KD, were more sensitive to apoptotic triggers

Because mitochondrial activity contributes to multiple cellular functions including ROS levels and sensitivity to apoptotic triggers, we wanted to determine how these processes changed upon modulation of TbSETD expression levels. Cells were incubated with ROS Assay Stain (Invitrogen) and exposed to peroxide for one hour (**Fig. 7**). In PF grown in 5 μM or 5 mM glucose media and BF parasites, we found significantly higher ROS levels in response to peroxide in TbSETD.HA.KD cells than in parental cell lines. In 5 μM glucose media TbSETD.HA.KD produced an average of ∼2.4 times more ROS than parental 29-13 (Mean fluorescence intensity (MFI) of 21,4687 ± 1880 in TbSETD.HA.KD and 87,024 ± 2851 in 29-13 cells) (**Fig. 7A**). In 5 mM glucose media TbSETD.HA.KD produced ∼ 2 times more ROS than 29-13 cells (MFI of 5814 ± 357 and 2915 ± 315 in TbSETD.HA.KD and 29-13 cells, respectively) (**Fig. 7B**). BF TbSETD RNAi cells (MFI = 821 ± 20) produced on average ∼ 1.4 times more total ROS compared to parental cells (MFI 575 ± 9.5) (**Fig. 7C**).

**Figure 7:**
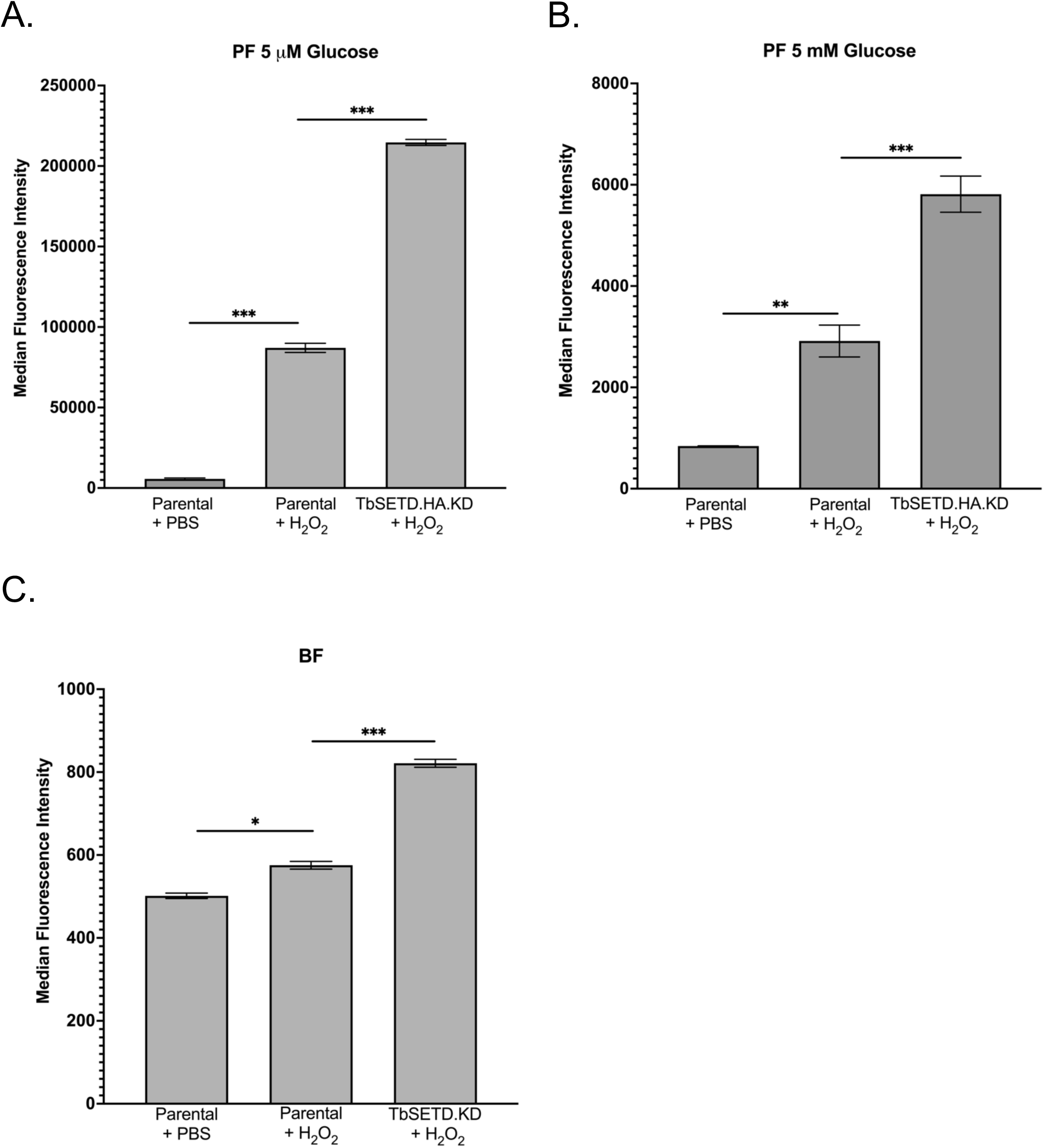
TbSETD.HA.KD cells show increased ROS levels. **A.** Bar graphs of relative ROS levels (median fluorescence intensity) of PF grown in 5 μM glucose **B.** Bar graphs of relative ROS levels (median fluorescence intensity) of PF grown in 5 mM glucose BF **C.** Bar graphs of relative ROS levels (median fluorescence intensity) of BF. Welch’s T-tests were performed (*P < 0.05, **P < 0.01, ***P < 0.001). All assays were completed in 3 biological replicates with 3 technical triplicates. Shown is one representative biological replicate; trends were the same across replicates.

In addition to influencing ROS levels, mitochondria are involved in mediating apoptotic events. While *T. brucei* lacks many proteins associated with apoptosis, they can initiate an apoptotic-like response to stresses like ROS, cytokines, and drugs (26). We induced this pathway with peroxide treatment and stained cells with annexin V-FITC, which binds phosphatidyl serine that is translocated from the inner to the outer leaflet of the plasma membrane during apoptosis and apoptotic-like responses (**Fig. 8**). PF TbSETD.KD cells had ∼25% and 33% more annexin positive cells in 5 μM glucose media and 5 mM glucose, respectively, than parental 29-13 (**Fig. 8A-B**). Additionally, ROS levels of both peroxide treated 29-13 and TbSETD3.HA.KD were higher in low glucose as compared to high glucose. Interestingly, in BFs, there was no difference in annexin V-FITC staining between parental and BF TbSETD.KD cells (**Fig. 8C**).

**Figure 8.**
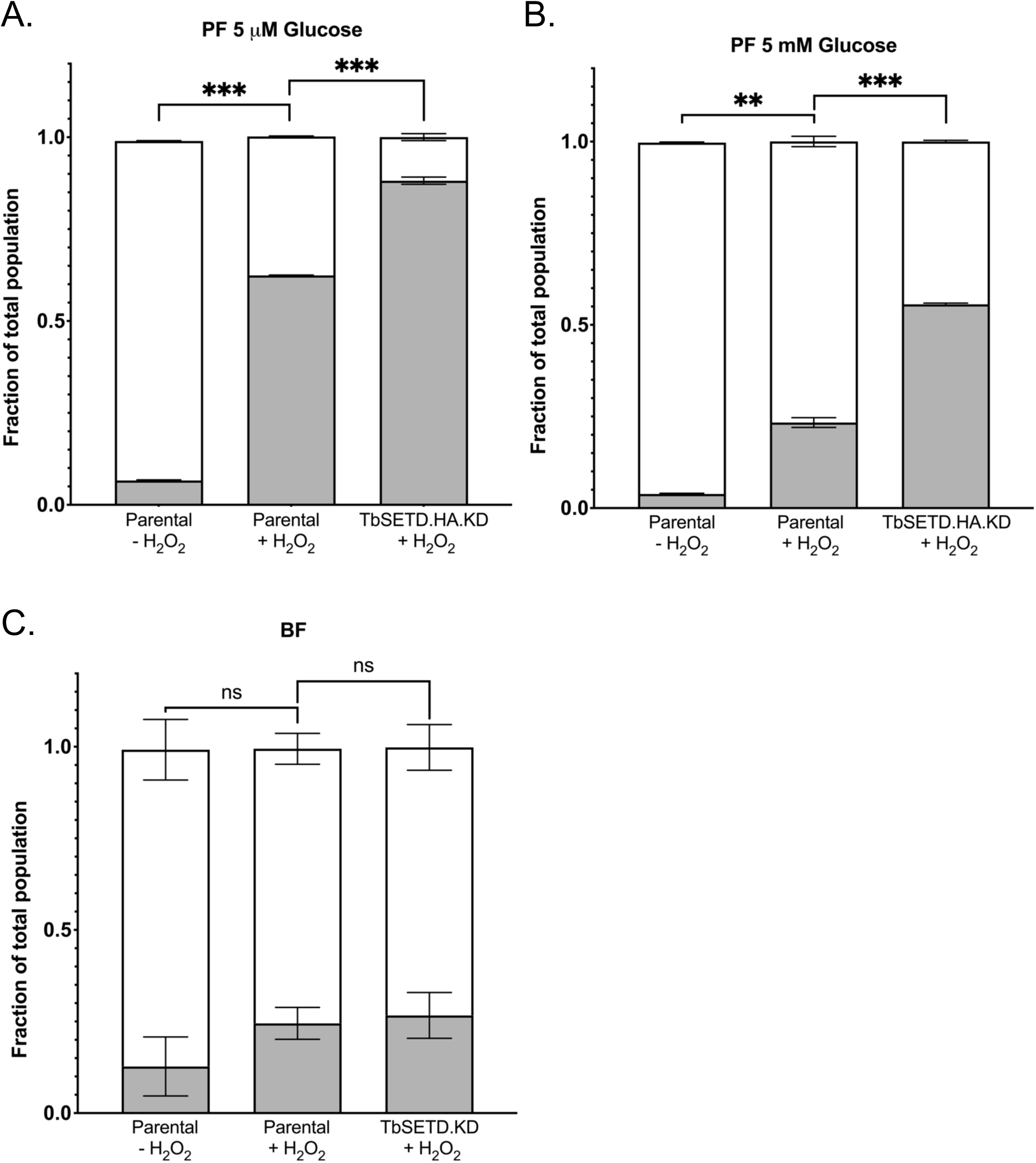
PF TbSETD.HA.KD exhibit increased sensitivity to apoptotic triggers. Bar graphs of annexin-positive cells in 29-13 and TbSETD.HA.KD cultures. **A.** PF cells grown in 5 μm **B.** PF cells grown in 5 mM glucose **C.** BF cells. Median fluorescence intensity was calculated, and Welch’s T-tests were performed (*P < 0.05, **P < 0.01, ***P < 0.001). All assays were completed in 3 biological replicates with 3 technical triplicates. Shown is one representative biological replicate; trends were the same across replicates.

### Potential TbSETD.HA binding proteins were enriched in mitoribosome and mitoribosome assembly proteins

To gain insight into potential TbSETD interacting proteins, we used mass spectrometry to identify proteins isolated by immunoprecipitations of TbSETD.HA from organelle-enriched fractions, which would include mitochondria (where a majority of TbSETD is localized) as well as glycosomes, acidocalcisomes, and ER microsomes. We reasoned that binding partners would include putative substrates as well as members of any larger protein complexes to which TbSETD might belong. Parental cell lysate was used to control for proteins that were isolated due to non-specific interactions with immunoprecipitation components. From the 99 proteins that were identified in the pulldown (**Fig. 9**, **Table 2**), 73 (∼ 62%) were annotated as having mitochondrial or kinetoplast localization. GO term analysis indicated an enrichment in structural constituent of ribosomes, gene expression, translation, mitochondria, and ribosome.

**Figure 9.**
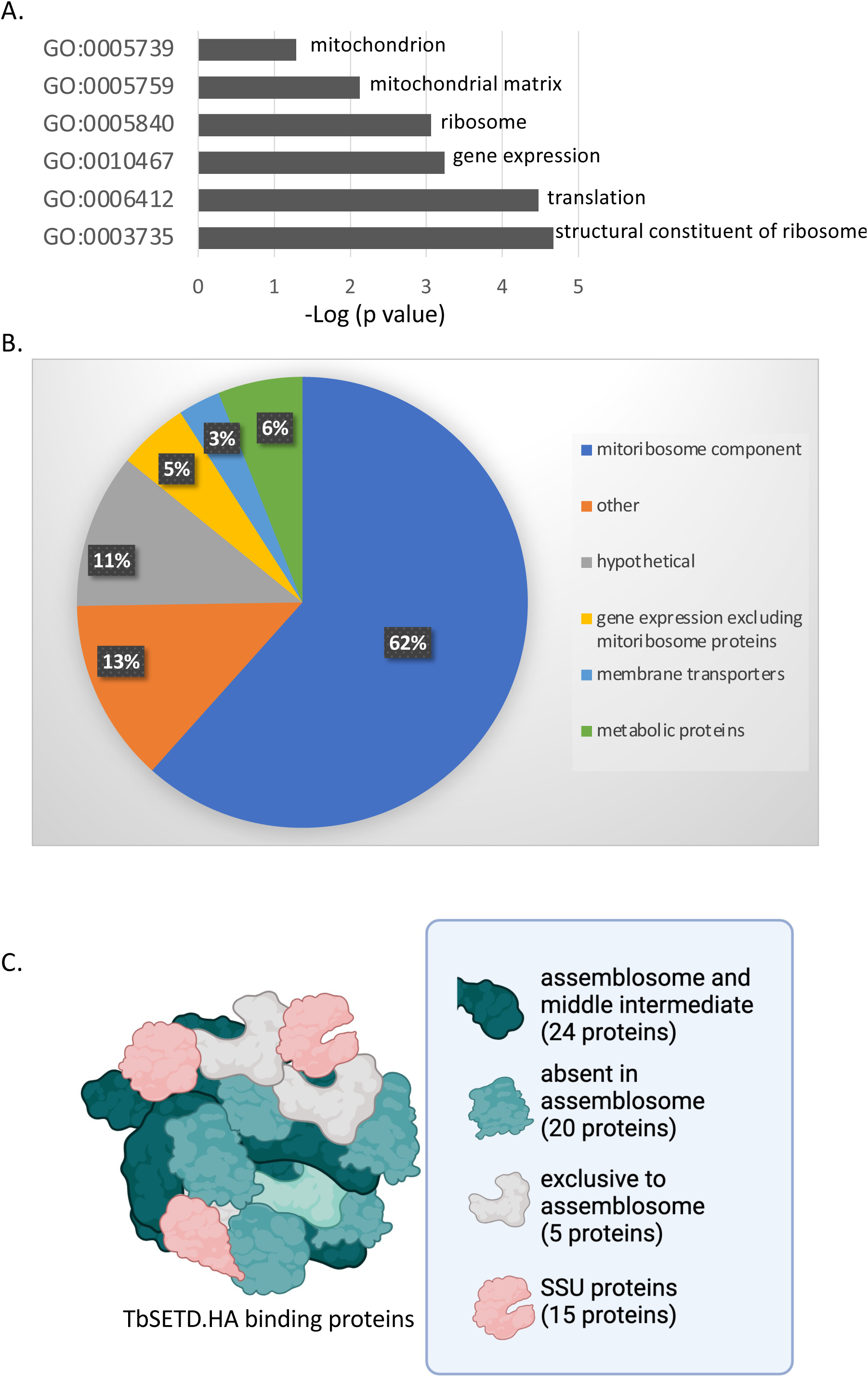
Mitoribosome assembly proteins are enriched in TbSET.HA immunoprecipitations. **A.** Bar graph of GO term enrichment analyzed via TriTrypDB.org. **B.** Pie chart of protein categories enriched in TbSETD.HA immunoprecipitations **C.** Mitoribosome small subunit assembly proteins present in TbSETD.HA immunoprecipitations. Colors indicate proteins present in specific mitoribosome assembly intermediates resolved previously (27). Dark green: proteins that were present in both early assemblosome and middle intermediate. Light green: proteins absent in early assemblosome but present in later intermediates. White: proteins exclusive to the assemblosome. Pink: mitoribosome small subunit proteins.

**Table 2.**
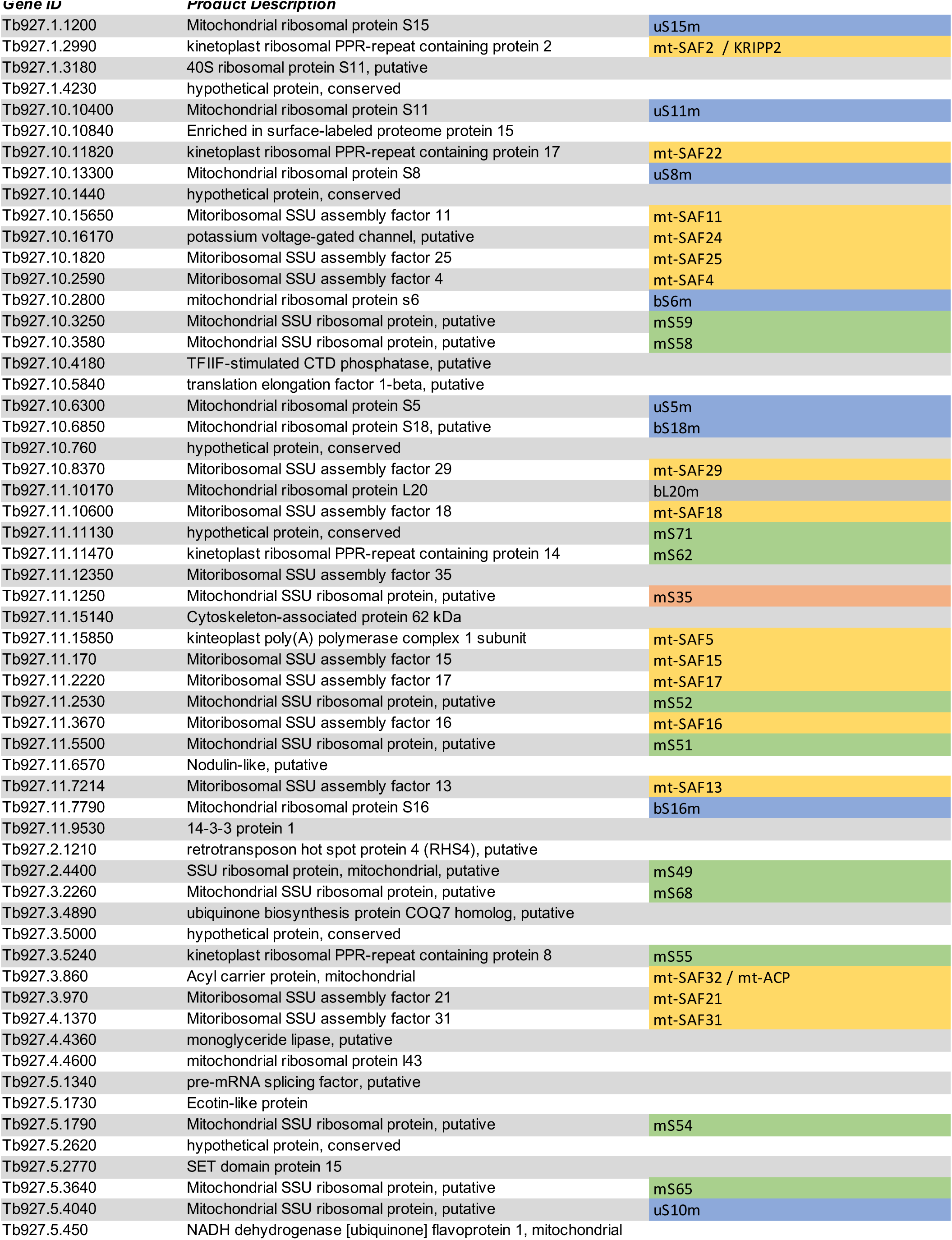
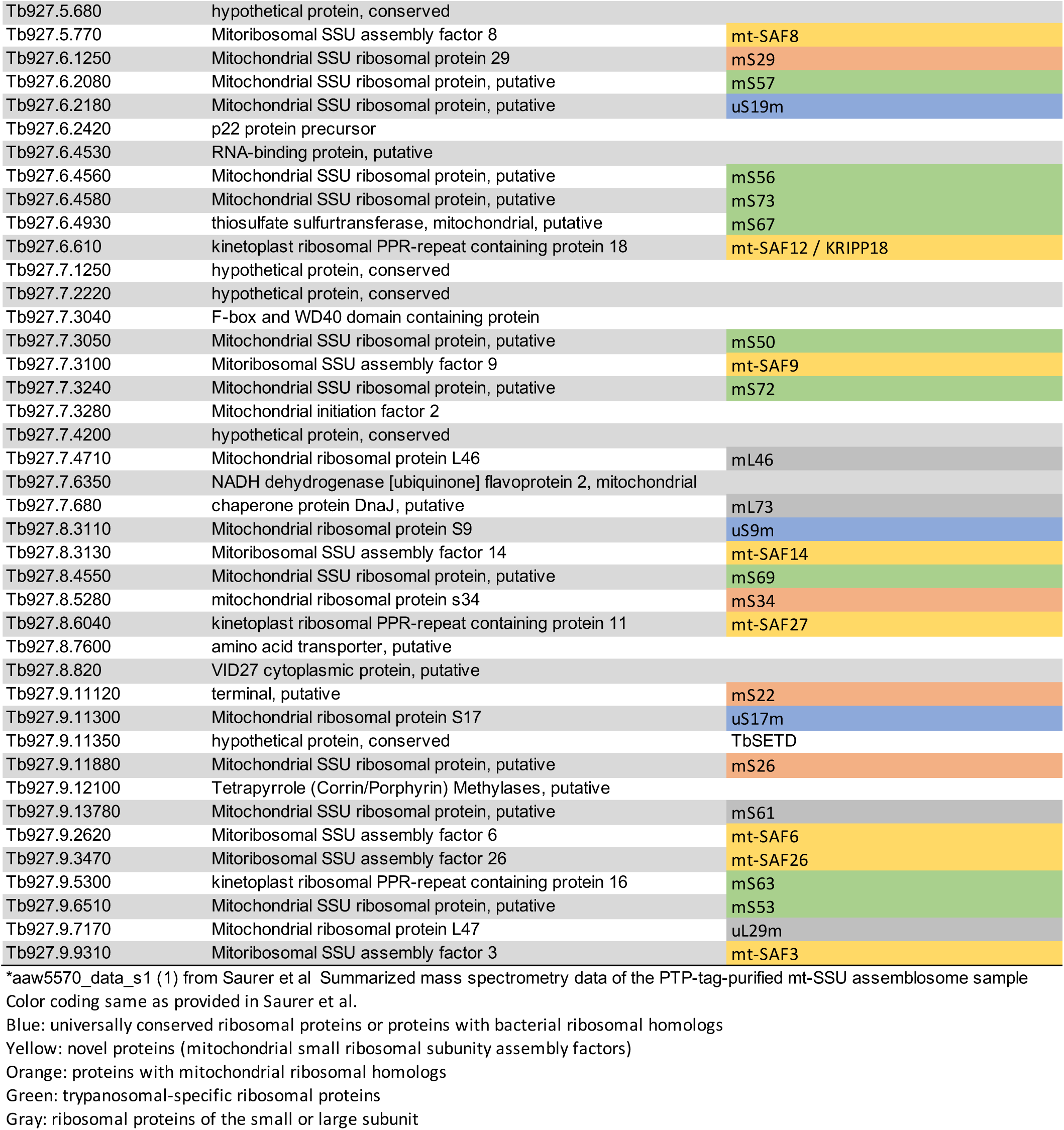
TbSETD binding proteins

Sixty-one proteins were mitoribosome proteins or mitoribosomal SSU assembly proteins. It was notable that 74 of the proteins we identified in our immunoprecipitations were also identified in PTP-tag-purified mt-SSU assemblosome (27). While the conditions used in our immunoprecipitations did not allow us to assess methylation, nine of the proteins identified in our pulldowns were methylated in *T. cruzi* epimastigotes. Most of these shared proteins were involved in translation (**Table 2, bold italics**) (28).

### TbSETD.HA purified from PF parasites exhibits methyltransferase activity and methylation of a 37 kDa protein is reduced in TbSETD.HA.KD cells

While TbSETD was recently annotated in TritrypDB as a SET domain protein, methyltransferase activity has not been documented. To address this gap, we tested whether TbSETD.HA purified from PF parasites had activity *in vitro*. We purified TbSETD.HA from parasites using magnetic anti-HA beads and resolved equal cell equivalents via SDS-PAGE and either stained with Coomassie or transferred to a membrane probed with anti-HA antibodies (**Fig. 10A**). PF 29-13 cells were used to control for activity from cellular components unrelated to TbSETD.HA. PF 29-13 TbSETD.HA samples contained a band at ∼ 34 kDa, the predicted size of the protein, which was absent from 29-13 parasites. We measured methyltransferase activity using a bioluminescence assay that scored the conversion of the methyl-donor S-adenosyl methionine to S-adenosyl-homocysteine. Activity was dose dependent and IPs from PF TbSETD.HA had ∼2X more activity (6,141 and 11,296 RLU) than 29-13 (2,850 and 6,750 RLU) (**Fig. 10B**).

**Figure 10.**
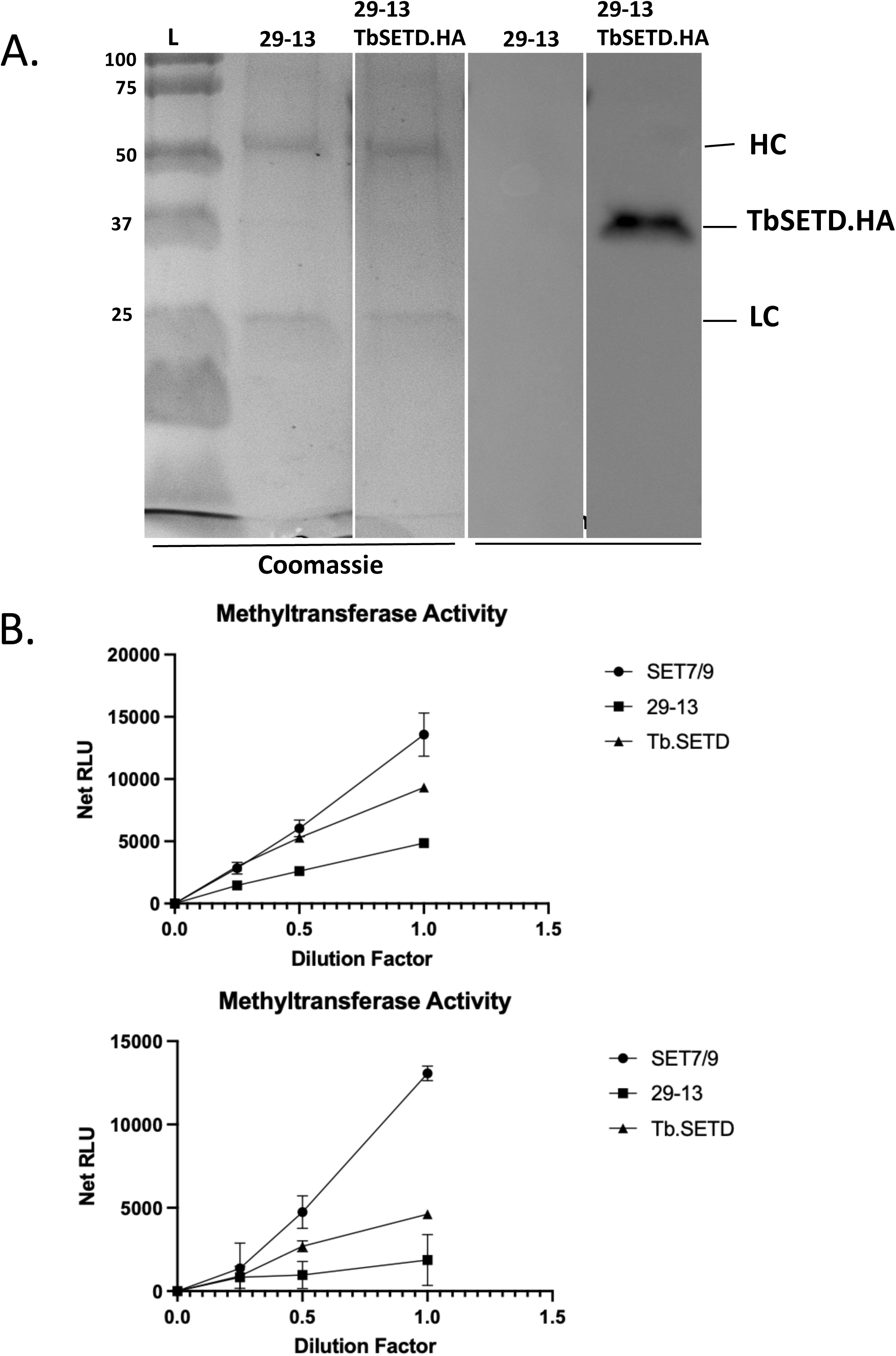
TbSET.HA IPs have methyltransferase activity. **A.** Coomassie-stained gel and western blot of immunoprecipitations from 29-13 or 29-13 TbSETD.HA probed with anti-HA. HC; heavy chain LC; light chain **B:** Methyltransferase activity of control human SET7/9 and immunoprecipitations fractions from parental 29-13 and 29-13 TbSETD.HA. Two biological replicates are presented with each assay done in triplicate. RLU; relative luciferase activity.

Because TbSETD is annotated as a protein lysine methyltransferase, and we detected methyltransferase activity in TbSETD.HA immunoprecipitations, we wanted to determine whether we could detect changes in protein lysine methylation in TbSETD.HA.KD cells. We induced silencing for 24 and 49 hour and probed mitochondrial fractions by western blotting with pan methyllysine antibodies. We observed an ∼ 75% reduction in the intensity of a 37 kDa protein (**Fig. 11**).

**Figure 11.**
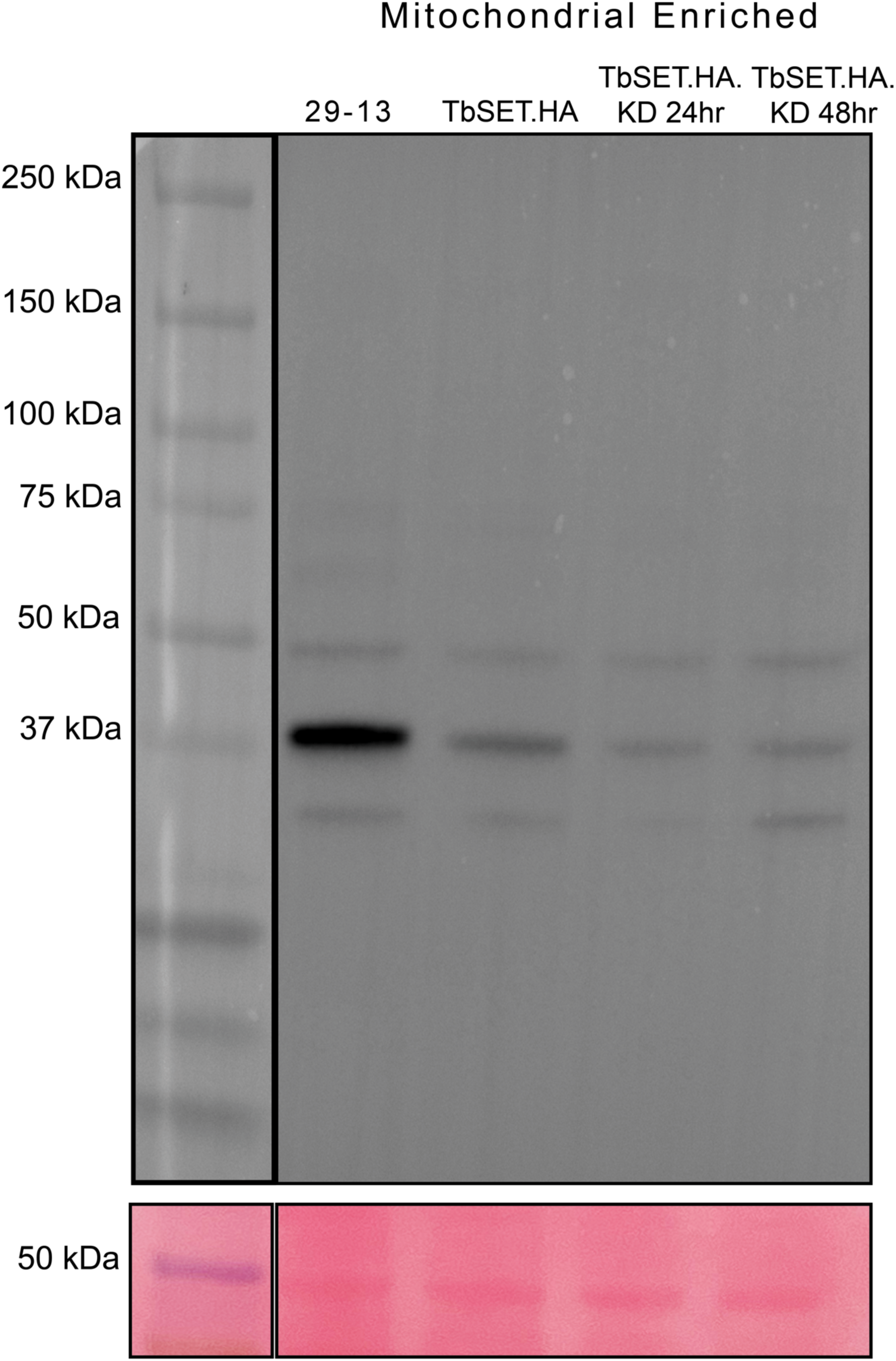
Methylation of a 37kDa band is reduced in TbSETD.HA.KD cells. Mitochondrial enriched fractions isolated from PF 29-13, TbSETD.HA, and TbSET.HA.KD induced for 24 h and 48h were resolved via SDS-PAGE, transferred to nitrocellulose membrane, and probed with pan-anti methylysine antibodies.

## Discussion

The major conclusions of the current study are that TbSETD is a novel SET domain protein that is essential in both PF and BF parasites and that, at least in PF, TbSETD is required for maintaining mitochondrial function and morphology, and interacts with many mitoribosome and mitoribosome assembly proteins. These conclusions are based on the following experimental evidence. First, we demonstrated that silencing TbSETD reduced growth in BF parasites and PF parasites grown in low glucose conditions. Next, we provided evidence that TbSETD localizes to the mitochondrion of PF parasites and that TbSETD-deficient cells had swollen mitochondria. Both PF and BF TbSETD-deficient cells has increased levels of ROS while only PF were more sensitive to apoptotic stressors. Finally, material from immunoprecipitations of epitope-tagged TbSET contained an exceptionally high number of mitoribosome and mitoribosome assembly factors and exhibited methyltransferase activity. Finally, methylation of a 37kDa protein was reduced in TbSETD.HA.KD cells. These findings are the first demonstration of a SET domain protein that is essential for parasite survival, the first documented mitochondrial SET domain PKMT in any eukaryote, and the first indication that protein lysine methyltransferases may be involved in mitoribosome assembly in any organism.

Humans have 57 SET protein domain proteins (29) (30), yeast have 12 SET domain proteins (31) and 9 SET domain proteins have been identified in *Plasmodium* (32). Our searches of the *T. brucei* genome identified 40 putative SET domain-containing proteins. Bioinformatic analyses identified homologs for 30 of these proteins in other organisms (8). Interestingly, 10 of the SET domain proteins we identified are annotated as hypothetical conserved or conserved with unknown function, and these lack homologs outside of the kinetoplastids, suggesting that they may be restricted to these protists.

Most of the proteins immunoprecipitated with TbSETD were mitoribosome or mitoribosome assembly proteins. Mitoribosomes translate 13 mitochondrially encoded proteins involved in the OXPHOS pathway. Trypanosome mitoribosomes are unique in that they contain less RNA and significantly more protein than mitoribosomes in other organisms (33). In fact, trypanosome mitoribosomes are the largest mitoribosome structures ever resolved by cryoelectron microscopy. Mitoribosome structure and assembly is an active area of research with many unanswered questions. A recent study on the mitoribosome small subunit (mt-SSU) of *T. brucei* has resolved the structures of four assembly complex intermediates designated as the mt-SSU assemblosome, middle assembly intermediate, late assembly intermediate, and the mature mt-SSU (27). In that work, 34 assembly factors involved in the maturation of the mt-SSU were identified.

TbSETD-binding proteins identified in immunoprecipitations performed in our lab include 24 of those 34 assembly factors as well as other protein components of the mt-SSU (**Table 2**). The enrichment of mt-SSU proteins along with the presence of TbSET in mitoribosome purifications performed in other laboratories (27), leads us to hypothesize that TbSET is involved in mt-SSU formation and/or function and may be part of a mitoribosome assembly intermediate.

A defect in mitoribosome function would have several consequences that are consistent with the phenotypes observed in our TbSETD-deficient cells. Decreased translation of mitochondrially encoded proteins involved in OXPHOS would result in a reduced capacity to generate ATP under low glucose conditions. In high glucose conditions, trypanosomes generate ATP primarily via glycolysis (34–37) and it would be reasonable to assume a defect in OXPHOS pathway would be less pronounced in high glucose conditions. This is consistent with our observation that the growth defect of PF TbSETD.HA.KD parasites was greater in low glucose conditions. Regardless of the molecular mechanism behind TbSETD function, the glucose-dependent phenotype of TbSETD-deficient parasites underscores the importance of studying PF parasite biology in both high and low glucose conditions. Additionally, OXPHOS proteins are membrane-bound and reduced levels of these proteins could lead to compromised mitochondrial membrane integrity and mitochondrial function. Interestingly, MitoTracker Red continued to label trypanosome mitochondria suggesting there is some membrane potential that is maintained in the TbSETD.HA.KD cells.

While lysine methylation of proteins in *T. brucei* has not been well studied, the significance of methylation as an important post-translation modification in parasite biology has been well established by studies focused on the methylation of arginine residues. About 10-15% of proteins are methylated at arginine residues in *T. brucei* and protein arginine methyltransferases influence glycolytic enzyme abundance, expression of proteins involved in proline degradation, messenger RNA stability and formation of mRNA granules upon stress (4, 38–40). In contrast to our knowledge base regarding arginine methylation, the role of lysine methylation, especially in relation to non-histone substrates, is unknown but likely plays an equally important role in parasite biology. We anticipate that lysine methylation and the mechanisms by which it regulates cell biology may share commonalities with arginine methylation.

The reduced abundance of a 37 kDa protein in methyllysine western blots of TbSETD-deficient cells may result from decreased protein expression, a decrease in methylation of the protein, or both. Additionally, we cannot eliminate the possibility that the methyltransferase activity observed in pulldowns of TbSETD is not a consequence of other proteins associated with TbSETD. We have tried unsuccessfully to express a variant of TbSETD in which the putative catalytic residue (Y268) corresponding to tyrosine Y261 in HsSETD is mutated to alanine. We have obtained drug resistant cell lines after transfection with expression constructs. However, we were unable to detect expression of epitope-tagged dead enzyme by western analysis or immunofluorescence assays. We propose that expression of this catalytically inactive protein is lethal, and that cell survival requires uncoupling of drug resistance from transgene expression.

Many SET domain proteins have multiple substrates. In addition to mitoribosome and mitoribosome assembly proteins, we also identified mitochondrial metabolic proteins in our immunoprecipitations including an acyl carrier protein (ACP, Tb927.3.860), NADH dehydrogenase [ubiquinone] flavoprotein 1 (Tb927.5.450), a putative thiosulfate sulfur transferase (Tb927.6.4930), a NADH dehydrogenase [ubiquinone] flavoprotein 2, (Tb927.7.63500), putative potassium voltage-gated channel (Tb927.10.16170) and a putative amino acid transporter, (Tb927.8.7600). Some of these metabolic enzymes, including ACP and the putative potassium voltage gated channel are documented members of mitoribosomes (41, 42). It is unclear whether the additional metabolic proteins identified in our immunoprecipitations are true TbSETD binding proteins or methyltransferase substrates. There is precedent for regulation of metabolism by lysine methyltransferases that belong to a different class of enzymes. In humans and yeast, there are mitochondrial methyltransferases belonging to the seven β-strand family, METTL20 and METTL12, that modify metabolic enzymes such as the β-subunit of electron transfer flavoprotein (43) and mitochondrial citrate synthase (44), respectively. Further work to define how protein expression, methylation, and metabolic activities are altered in the TbSETD.HA.KD cells is required resolve the molecular mechanisms behind the phenotypes we observed in our studies. To our knowledge, there are no published studies on SET domain mitochondrial methyltransferases.

In PF parasites, V5.TbSETD localized to the mitochondria. We should note that we could not express epitope-tagged TbSET in BF, so we do not know if the protein localizes to the mitochondrion in both lifestages. In addition to TbSETD, four of the TbSET proteins annotated as hypothetical or conserved unknown were detected in the mitochondrion of parasites in TrypTag project (45, 46)(**Table 1**), suggesting there may be additional SET proteins involved in mitochondrial function.

Initial silencing experiments were performed in cells expressing TbSETD.HA so that we could monitor the penetrance of RNAi as we don’t have antibodies against the native protein. It is possible that the phenotype observed in these cell lines is a consequence of both expression of a tagged TbSETD as well as a reduction in overall TbSETD levels. While we can monitor expression of tagged TbSET, our lack of antibodies prevents us from measuring levels of wild-type protein. To uncouple these two effects, we measured growth rates of cells expressing TbSETD.HA lacking the RNA interference plasmid and cell transformed with the silencing construct alone. Both cell lines exhibited growth defects (supplementary figure 2 and 3) indicating that both expression of TbSETD.HA and silencing of TbSETD is detrimental to parasite growth. Long-term experiments were performed with the TbSETD.HA.KD cell lines because cells harboring the silencing construct alone do not survive cryostorage and must be transformed prior to each experiment.

In summary, we show here than TbSETD, an unusual SET domain protein that is conserved in kinetoplastids and absent in other eukaryotes is essential in both PF and BF parasites and required for proper mitochondrial function. While protein lysine methylation of non-histone proteins serves a fundamental role in eukaryotic cell biology, this is the first mitochondrial SET domain protein characterized in *T. brucei*. Furthermore, it is the first implication of a protein methyltransferase that may be associated with mitoribosome assembly and/or function. Future studies will be focused on identifying TbSETD substrates *in vivo* and resolving the molecular mechanism by which TbSETD regulates mitochondrial function.

## Acknowledgments

We would like to thank Wandy Beatty (Washington University, Molecular Microbiology Imaging Facility) for the electron microscopy images and Drs. Sam Mackintosh and Stephanie Byrum (University of Arkansas for Medical Sciences, Mass spectrometry facility) for the mass spectrometry data.

## Supplemental Figures

**Supplemental figure 1.**
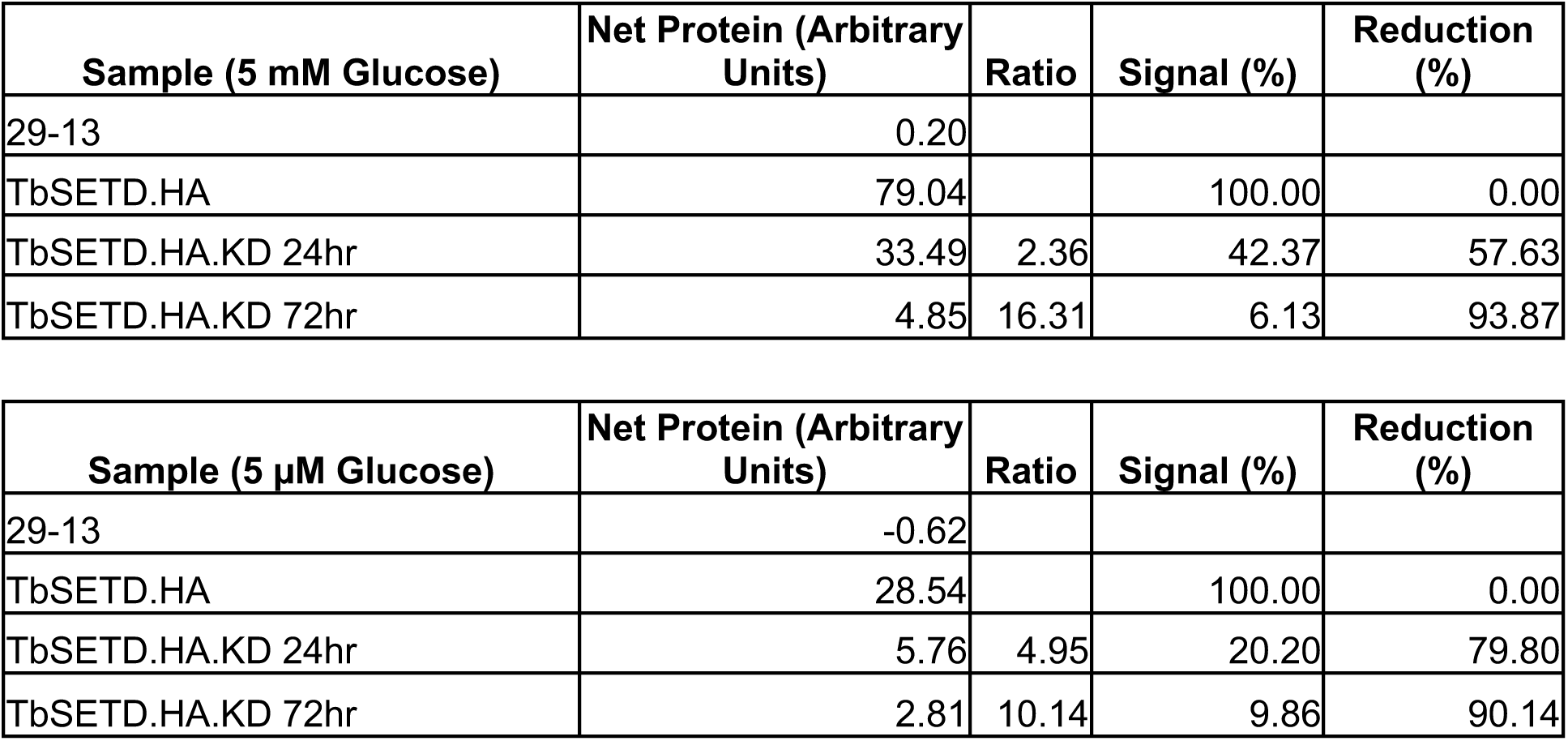
Quantification of TbSETD.HA silencing. Bands in western blots in Fig. 3 were quantified via densitometry using Fiji and reported as Net Protein (Arbitrary Densitometry Units). Ratio is fold-change of the silenced TbSETD.HA Net Protein in comparison to TbSETD.HA. TbSETD.HA expression levels were set as 100%. Signal % is equal to the amount of signal compared to TbSETD.HA and percent reduction is equal to the amount of protein lost due to RNAi. FIJI (47): No plug-ins used

**Supplemental figure 2.**
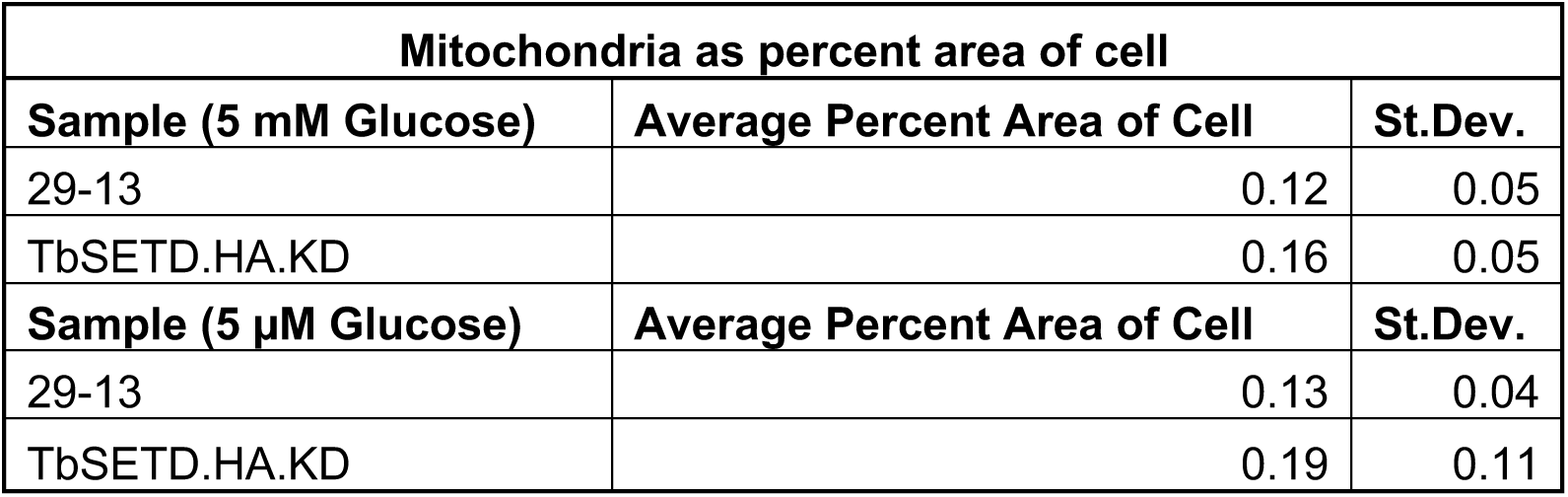
Values for mitochondrial area. Mitochondria area was measured via FIJI (47) and calculated as a percentage of cell area.

**Supplemental Figure 3.**
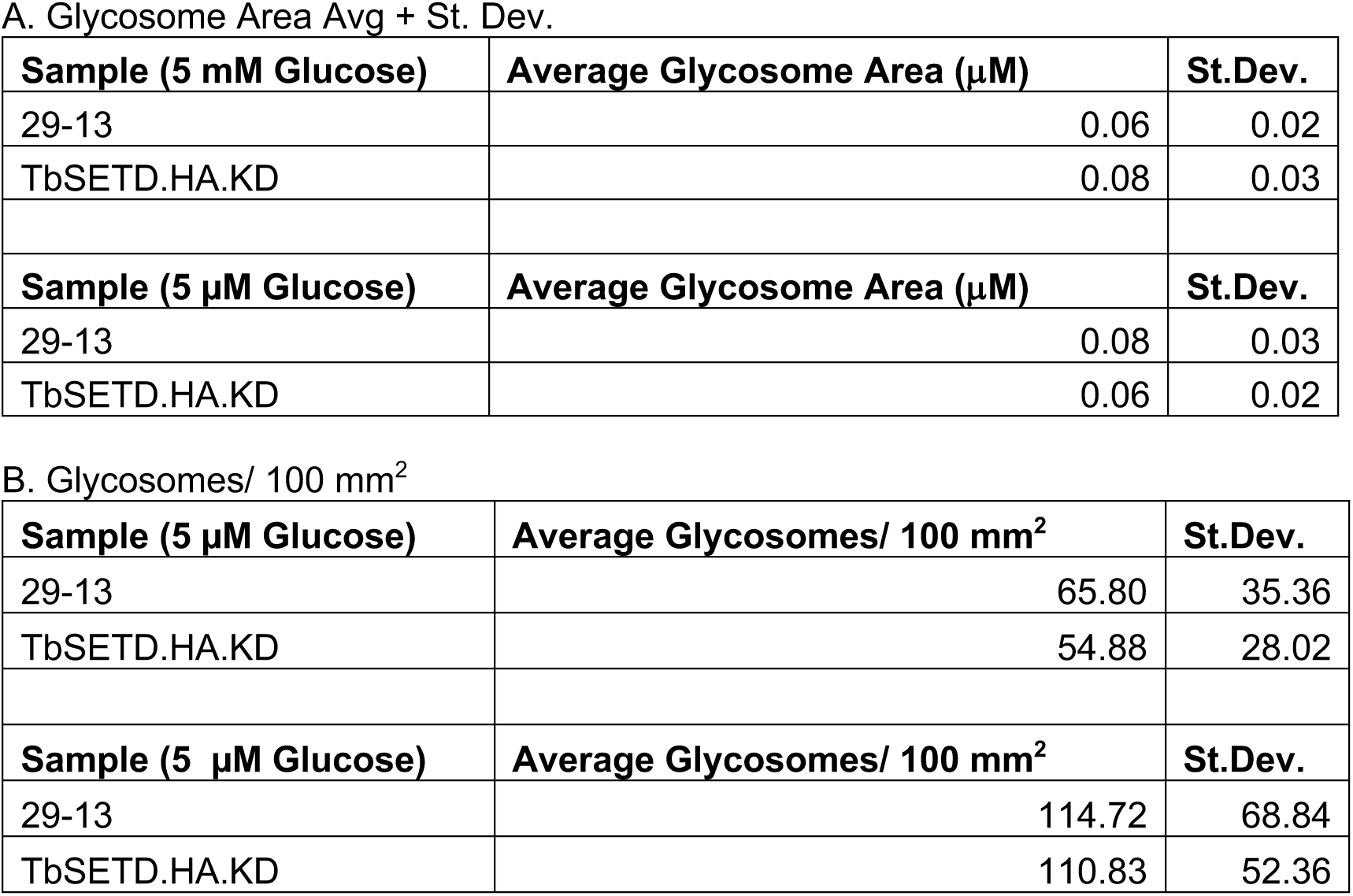
Values for glycosome number and area. Glycosome size and number were measured via FIJI (47) and calculated as a percentage of cell area. Values for glycosome number and area. Glycosome size and number were measured via FIJI (citation) and calculated as a percentage of cell area.

**Supplemental Figure 4.**
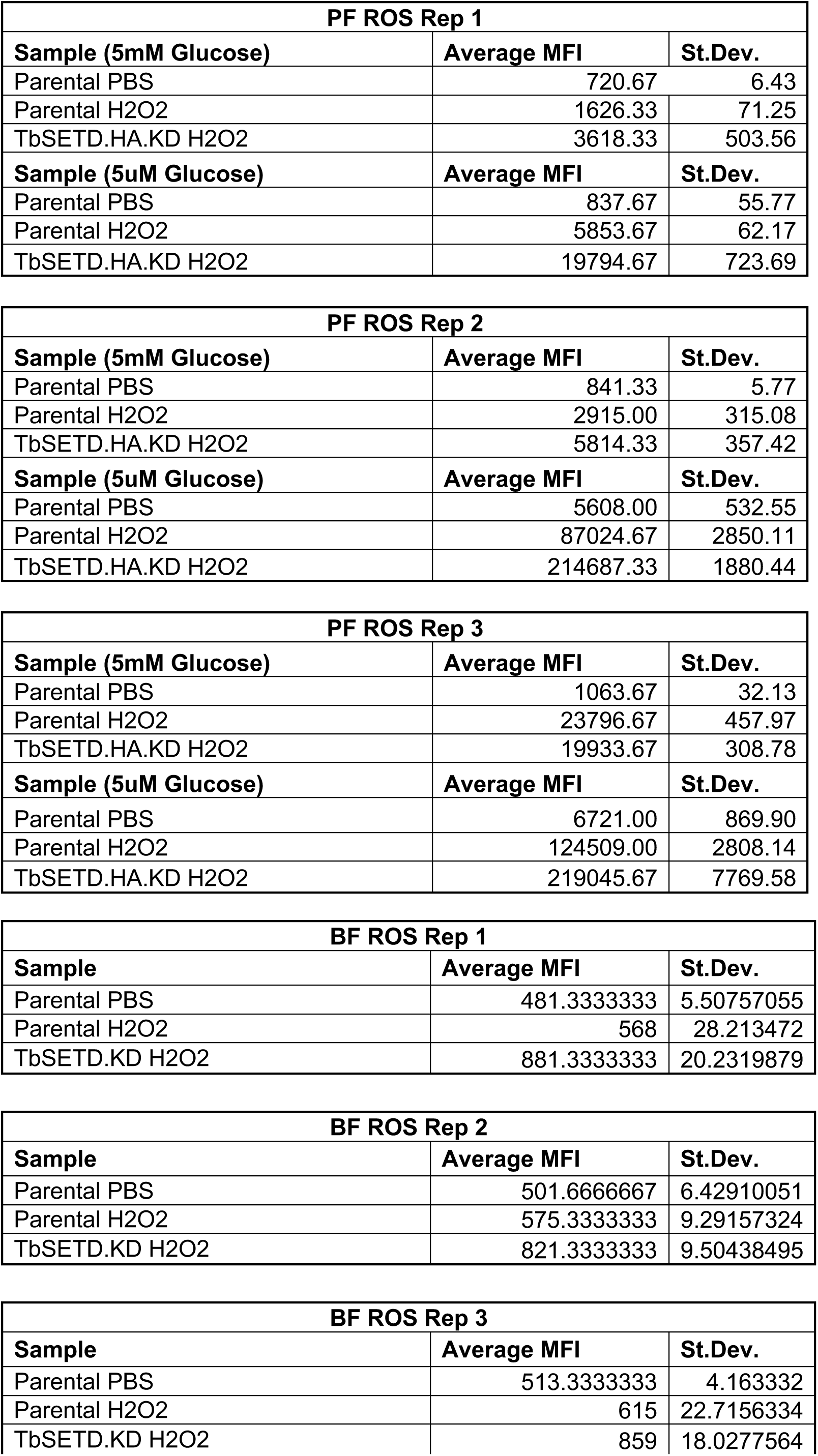
ROS assay values of mean fluorescence intensities (MFI). Data was recorded for three biological replicates, each with technical triplicates. Data from replicates 2 of PF and BF were used in Fig. 7.

**Supplemental Figure 5.**
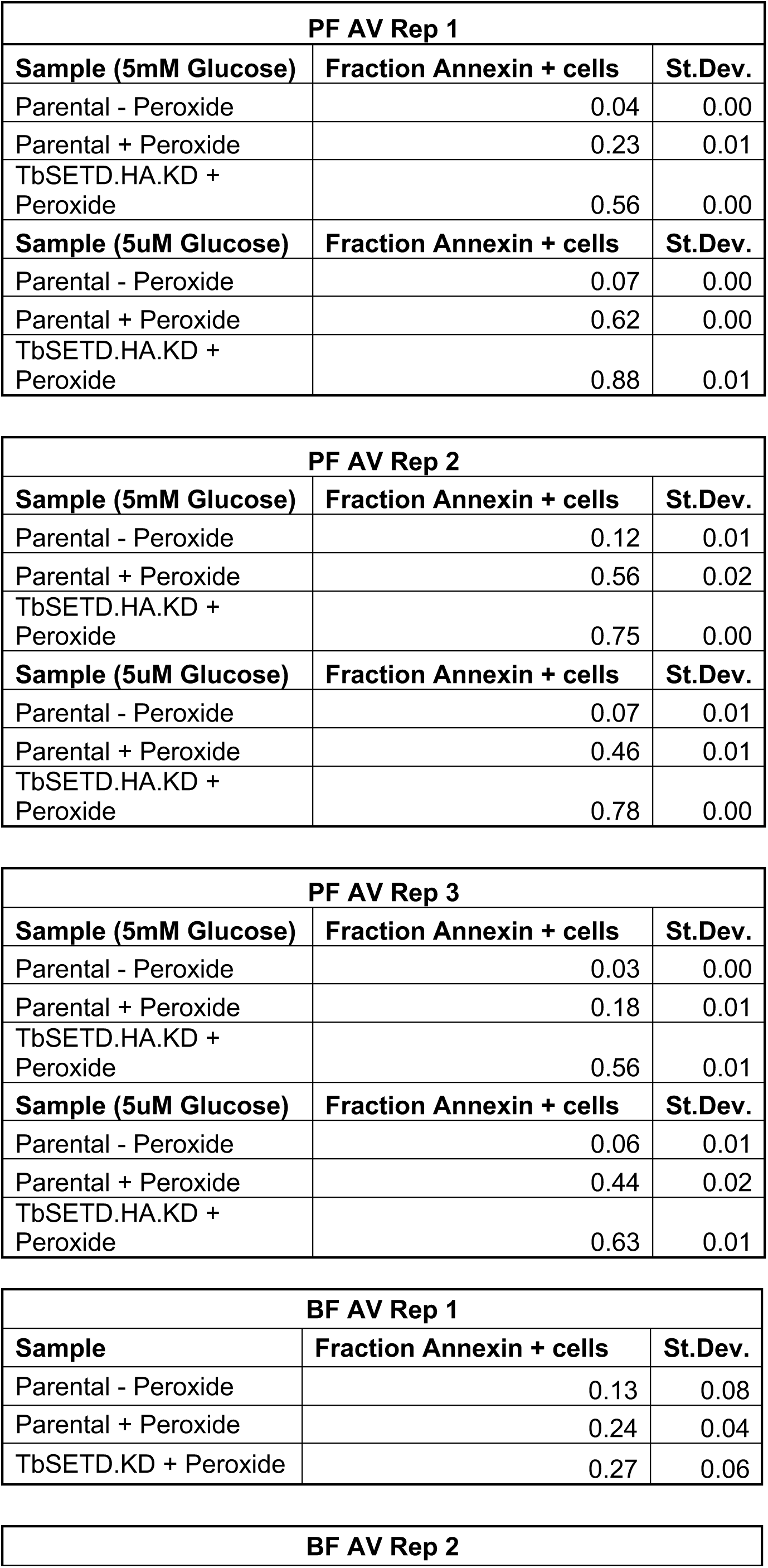
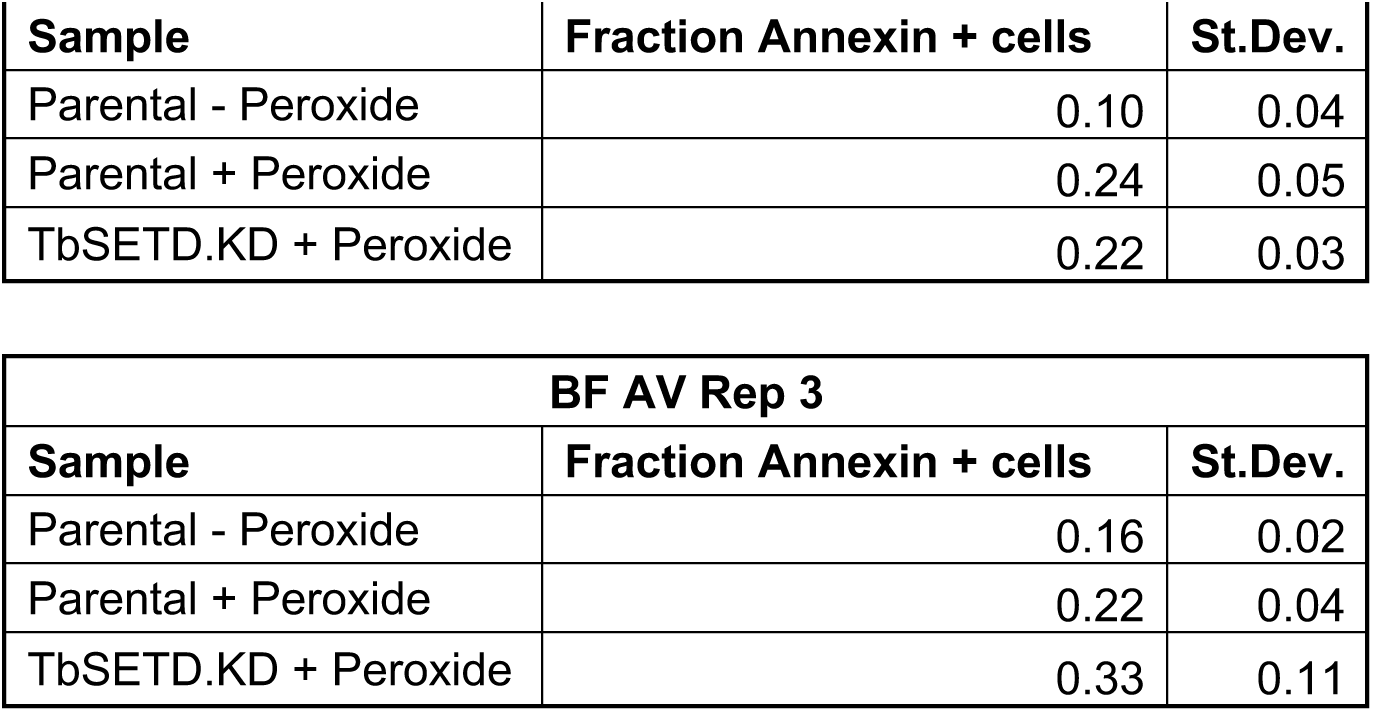
Annexin V positive cells reported as the fraction of total population. Each experiment consisted of three biological replicates, each with technical triplicates. Data from replicates 2 of PF and BF were used in Fig. 7

**Supplemental Figure 6.**
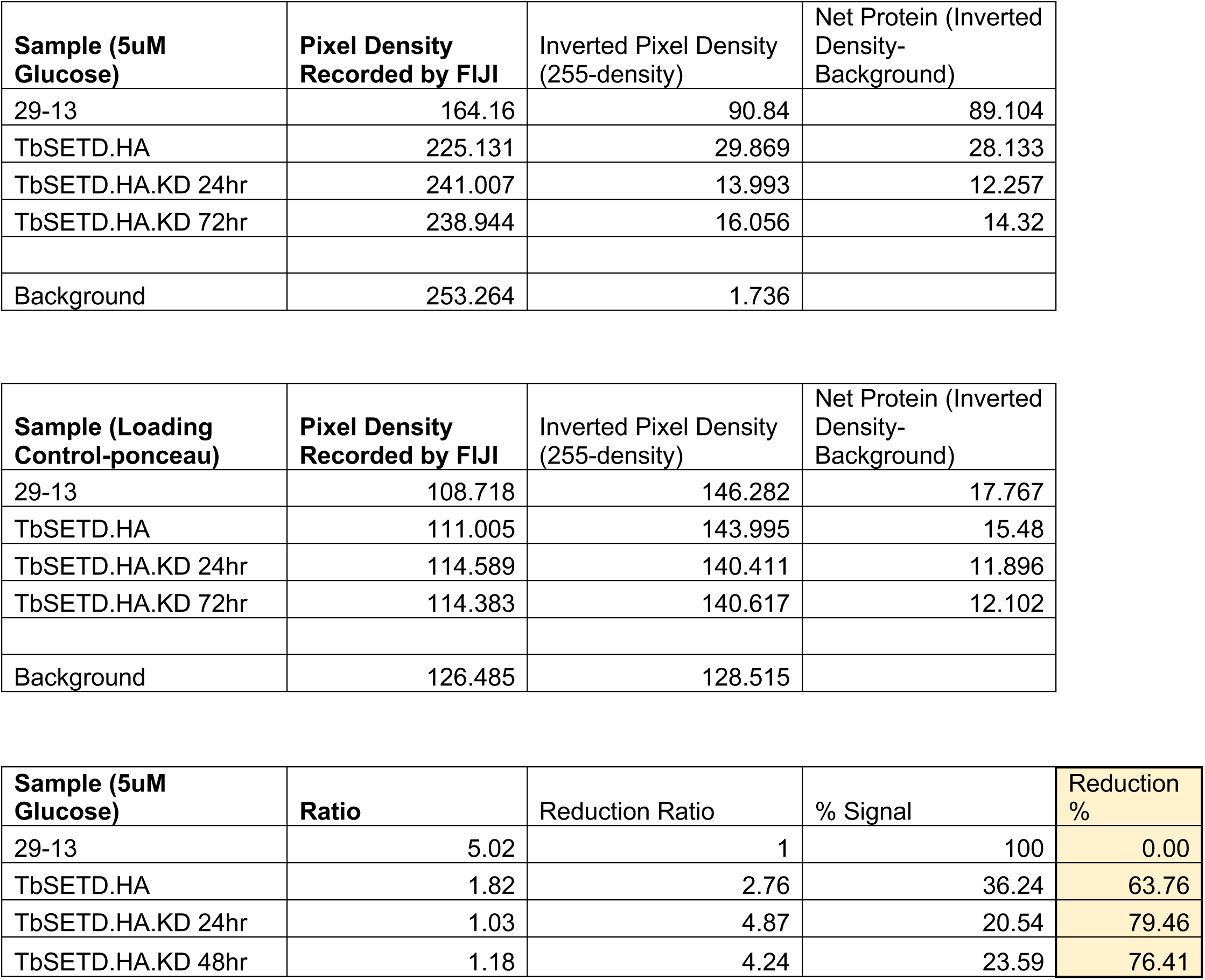
Densitometry values for methyllysine blots. 37 kDa bands in Western blot in Fig. 11 were quantified via densitometry using Fiji and reported as Net Protein (Arbitrary Densitometry Units). Ratio is the Net Protein value over the Net Loading control for each sample. Reduction Ratio is fold-change of the modified TbSETD sample’s Net Protein in comparison to 29-13. 29-13 levels were set as 100%. Signal % is equal to the amount of signal compared to 29-13, and reduction % is equal to the amount of protein lost due to TbSETD modifications.

**Supplemental Figure 7.**
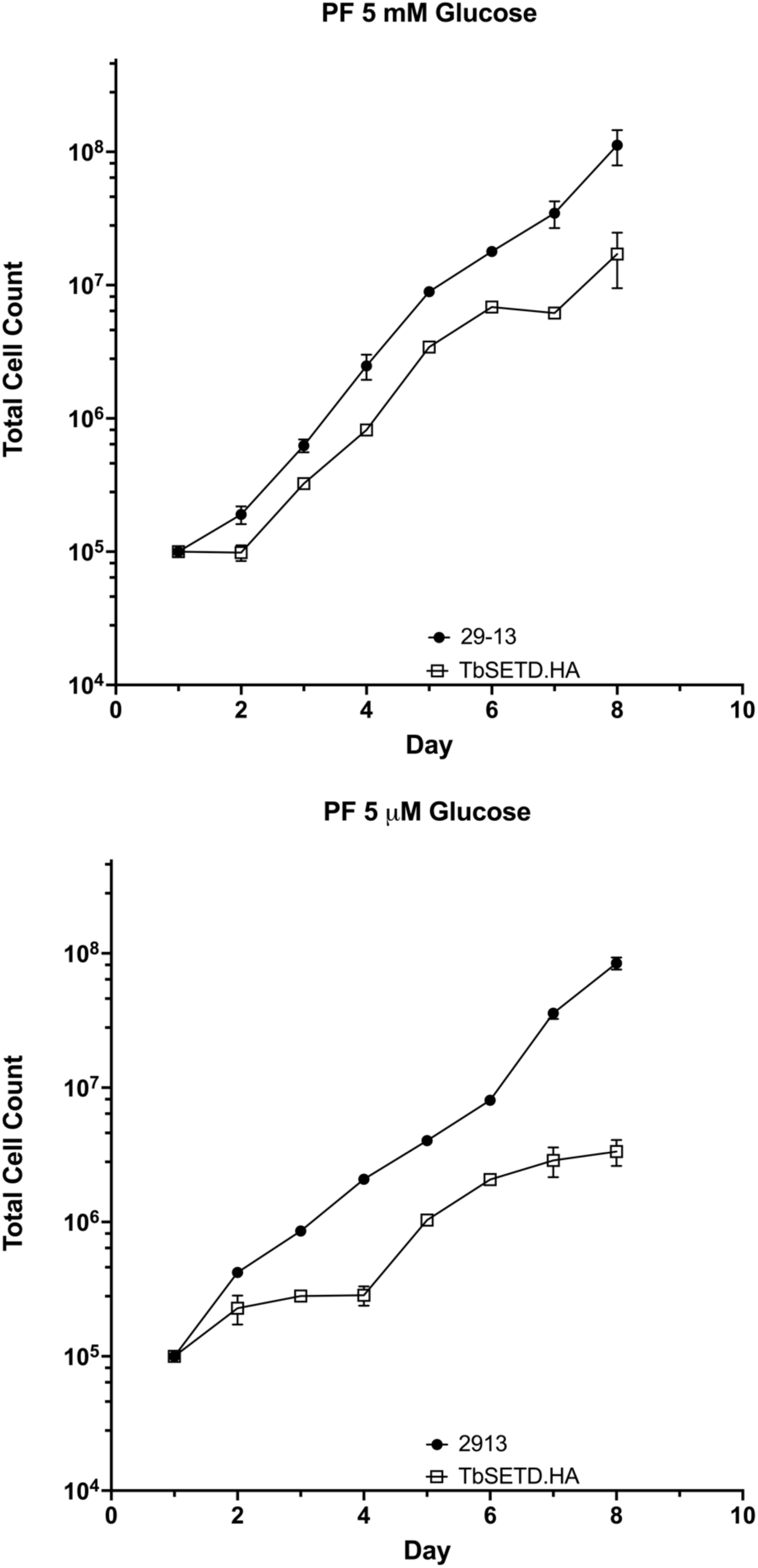
Growth curves of PF 29-13 TbSETD.HA. PF parasites expressing TbSETD.HA were counted daily. **A.** PF parasites grown in 5 mM glucose. **B.** PF parasites grown in 5 μM glucose.

**Supplemental Figure 8.**
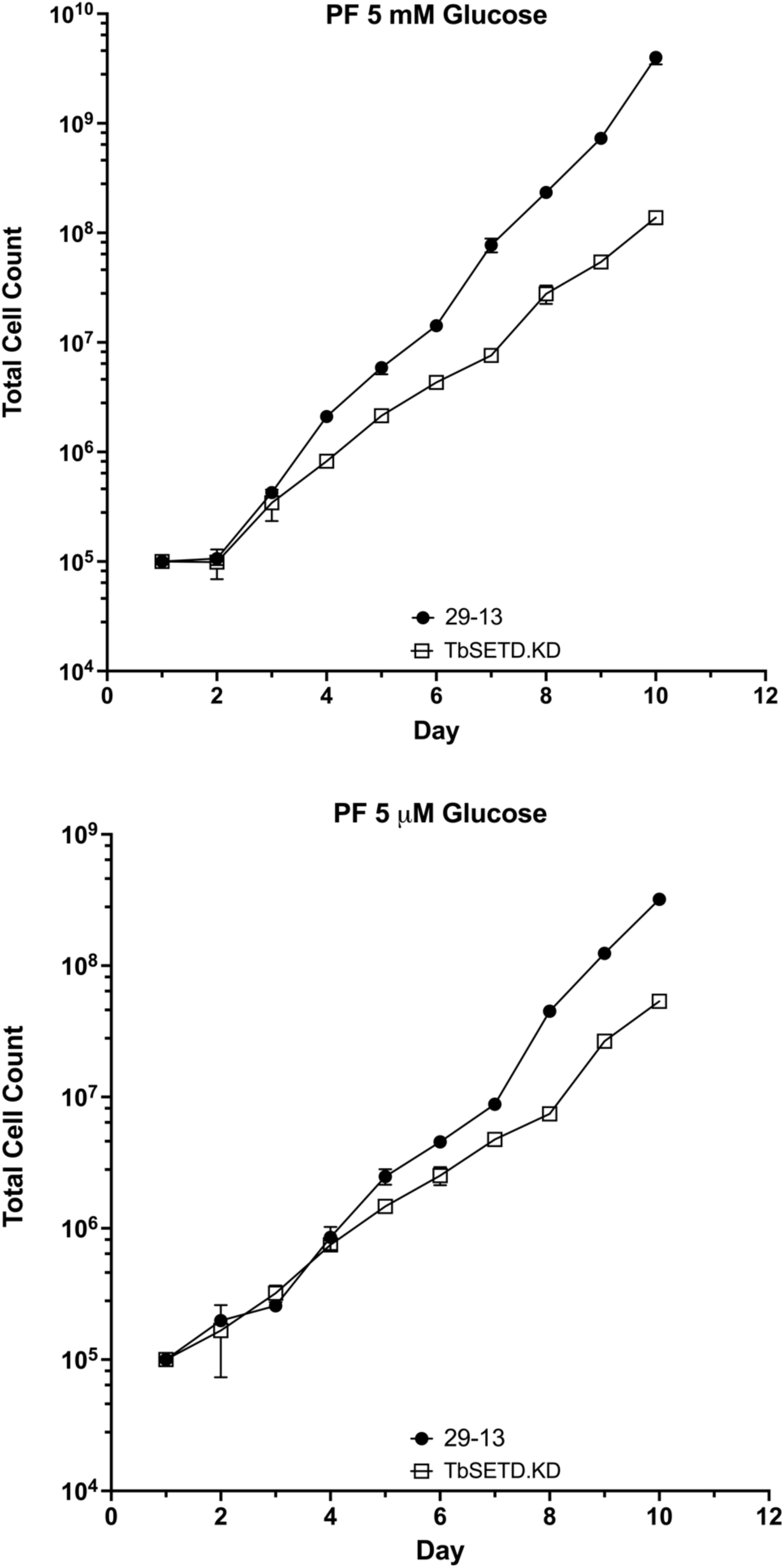
Growth curves of PF 29-13 TbSETD.KD. PF parasites harboring pZJMTbSETD construct for gene silencing were induced with doxycycline and counted daily. **A.** PF parasites grown in 5 mM glucose. **B.** PF parasites grown in 5 μM glucose.

**Supplemental Figure 9.**
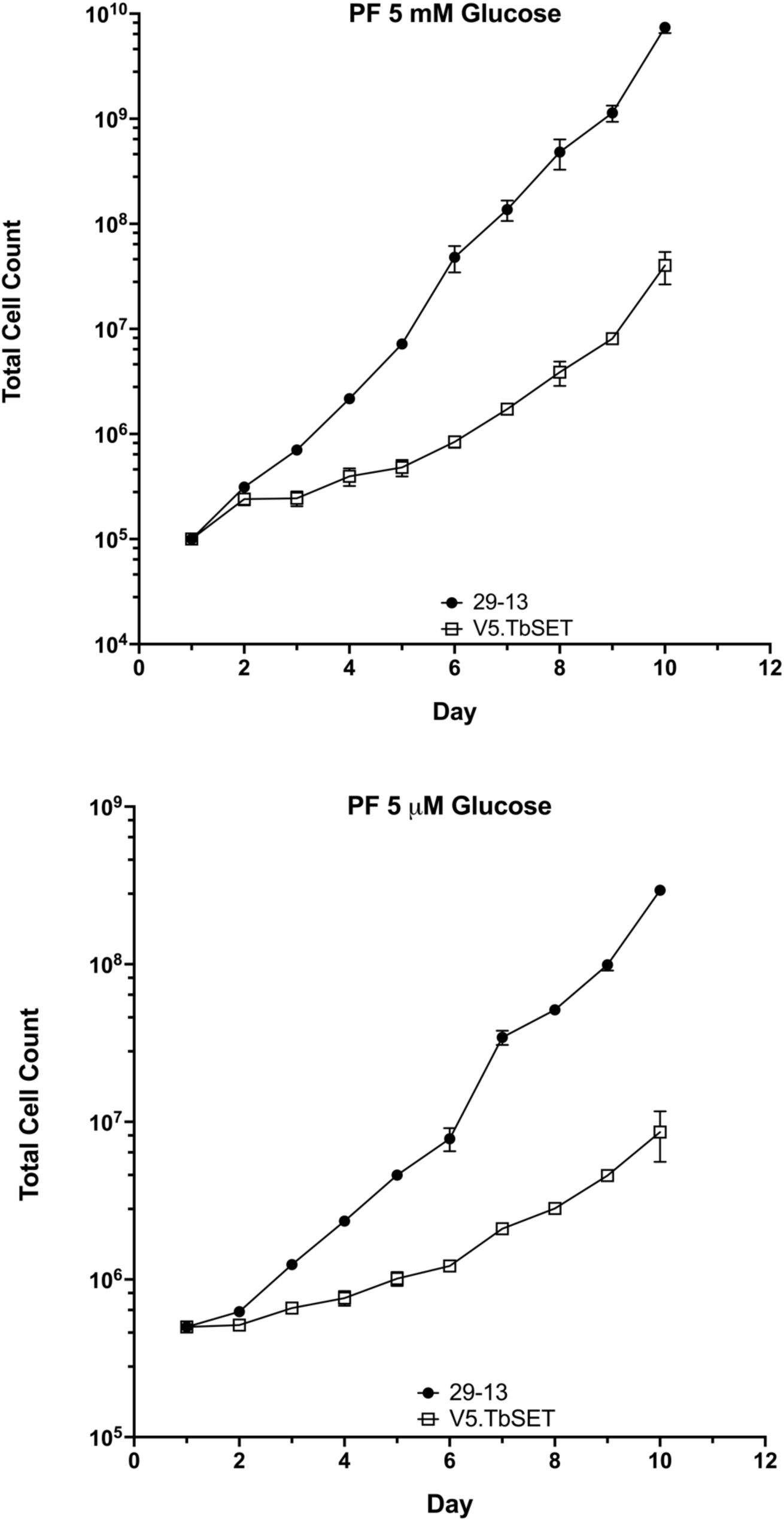
Growth curves of PF 29-13 V5.TbSETD. PF parasites expressing V5.TbSETD were counted daily. **A.** PF parasites grown in 5 mM glucose. **B.** PF parasites grown in 5 μM glucose.

